# Bioarchaeological analysis of one of the earliest Islamic burials in the Levant

**DOI:** 10.1101/2020.09.03.281261

**Authors:** Megha Srigyan, Héctor Bolívar, Irene Ureña, Jonathan Santana, Andrew Petersen, Eneko Iriarte, Mattias Jakobsson, Colin Smith, Juan José Ibañez, Anders Götherström, Torsten Günther, Cristina Valdiosera

## Abstract

The Middle East plays a central role in human history harbouring a vast diversity of ethnic, cultural and religious groups. However, much remains to be understood about past and present genomic diversity in this region. Here, we present for the first time, a multidisciplinary bioarchaeological analysis of two individuals dated to late 7th and early 8th centuries from Tell Qarassa, an open-air site in modern-day Syria. Radiocarbon dates, religious and cultural burial evidence indicate that this site represents one of the earliest Islamic Arab burials in the Levant during the Late Antiquity period. Interestingly, we found genomic similarity to a genotyped group of modern-day Bedouins and Saudi rather than to most neighbouring Levantine groups. This is highlighted through substantial Neolithic Levant ancestry in our samples, inviting an alternative scenario of long-term continuity in this region. This raises questions about the influence of ancient populations and historical migrations to genetic structure in the Middle East. As our study represents the first genomic analysis of an early Islamic burial in the Levant, we discuss our findings and possible historic scenarios in light of forces such as genetic drift and their possible interaction with religious and cultural processes.

## Introduction

The Middle East has an unparalleled place in human history. From the out-of-Africa movements of modern humans and admixture with Neandertals, to the spread of agriculture and emergence of civilisations, it has been at the crossroads of genetic as well as cultural history for millennia. While archaeological and historical sources provide instrumental insights into social/cultural aspects, ancient DNA (aDNA) enables the recovery of past genetic information, filling crucial gaps in our understanding of history. A number of studies on ancient DNA from different time periods in the Middle east have provided a general overview of the genetic history in this region. These include descriptions of the earliest local farming groups from the Neolithic^1^ and their expansions into Europe^2,3^ as well as the genetic differentiation among contemporary Neolithic groups ^4–6^. In the later Chalcolithic period, evidence of distinctive cultural practices and associated population movements highlight the dynamic history of the region, especially in the Southern Levant^7,8^. Further, genomic studies from the Bronze to Iron Ages in the Levant also report admixture and population movements, and suggest some degree of continuity with modern populations^6,9–11^. On an even more recent timescale, studies from the medieval period^6,11^ and modern populations^13,14^ describe genetic structure and the role played by culture and religion in the formation of these structures. Notably, a study on medieval individuals from Lebanon previously identified as Crusaders^12^ demonstrated their ancestry to be both European or local as well as an admixture of Europeans and Near-Easterners. These signals cannot be detected in modern Lebanese groups, suggesting these were only transient. This provided evidence of genetic signatures of historical religious events such as The Crusades which saw considerable movement from Europe to the Middle East, admixture with local populations and eventually, their dilution over time. Genetic analysis of modern Lebanese populations^13^ suggest that although ancient population movements in the Epipalaeolithic contributed to the divergence of Levantines from Europeans, the spread of Islam in the past millennium led to stronger stratification. Further south, data from modern Yemenis combined with other Middle Eastern groups found little correlation between genetic structure and geography^14^. It is thus clear that the present distribution of genetic diversity in the Middle East is the result of convoluted processes with culture as an additional level of complexity.

The Late Antiquity period, roughly defined as the time between the 3^rd^-8^th^ centuries, was a time of cultural and religious upheaval in the Middle East as well as in Europe. Byzantine Syria-Palestine was conquered by Islamic Arabs in the first half of the 7^th^ century AD (630s), becoming the political centre of the emerging Arab Islamic empire when the Umayyad caliphate was founded in Damascus in 661^15^. However, the Arabization and Islamization of the area did not fully take place until the last decade of the 7th century led by ‘Abd al-Malik^16^. As such, the Aramaean, Byzantine and Christian legacy interacted with the new Arab Islamic rule and cultural values for decades, accompanied by influx of populations from the Arabian Peninsula. The collapse of the Umayyad caliphate (750) and transfer of the political centre to Iraq caused political marginalization and economic decline in Syria-Palestine^17^ (see also Supplementary Text S1 – Early Islamic Southern Syria). Ancient DNA analysis is a powerful tool to provide a genomic snapshot of this dynamic period giving insight into past demographic processes of a currently conflicted and inaccessible territory of the Levant. Interestingly, despite the fact that much of the focus of Near Eastern Archaeology has been on funerary remains, surprisingly, few Islamic burials have been investigated. We present an archaeogenomic analysis of two Late Antiquity individuals buried with evidence of Islamic funerary rituals in modern-day Syria making their genomic study an invaluable resource to understand their ancestry and the history of the region.

## Results

### Description of the burials

In this study, we perform genomic analyses of two buried individuals excavated at Tell Qarassa North, a Neolithic site in the Village of Qarassa in Syria (Fig. 1a). While Tell Qarassa North is usually known as a prehistoric site^18,19^(see also Supplementary Text S1), the two individuals analysed here were found on surface levels of the site and directly radiocarbon dated to the Late Antiquity period (7^th^ to 8^th^ centuries) (Table 1). No cultural artifacts were associated with the human remains, however, their archaeological context revealed that these were primary burials with the bodies placed in decubitus position, inside pits that were intrusive in the Neolithic levels (Fig. 1b). Interestingly, while the two burials were located very close to each other, no evidence of a Late Antiquity cemetery was documented at the site. The individual syr005 was found laid on his back in a decubitus supine position, although the lower limbs were slightly flexed and placed on their right side. The burial was oriented east-west, with the head at the west, facing south. Further, individual syr013 was placed on her right side in a lateral decubitus position oriented east-west, with the head at the west, facing south. The distribution of the skeletal elements suggests that both bodies were wrapped before burial^20^. When soft tissues of the body decay faster than the wrapping, it can create either temporary or semi-permanent spaces around the putrefied body, yielding some skeletal movements at the disarticulated joints before wrapping decay (i.e. right elbow in individual syr013, left shoulder in individual syr005). The wrapping, the position and orientation of the bodies facing Mecca are concordant with Muslim funerary rituals following Early Islamic burials^21^, however, these individuals were not buried in a traditional Muslim cemetery. This may be explained due to special circumstances of death or cultural identity: nomadic populations, pilgrims, deviant burials or plague victims. The requirement of a Muslim burial to take place within 24 hours after death might have made some compromises necessary. It is known that one of the defining features of Muslim burials is that of only one person per grave, which implies that husbands and wives are not buried together, and collective family tombs are forbidden. However, occasionally and in extreme circumstances this can be relaxed for victims of plague or warfare (Supplementary Text S1-Muslim Burials). Also, the close proximity of radiocarbon dates for syr005 (1294 ± 18 Cal BP) and syr013 (1302 ± 15 Cal BP) suggest that both individuals died at a similar time.

**Fig. 1.**
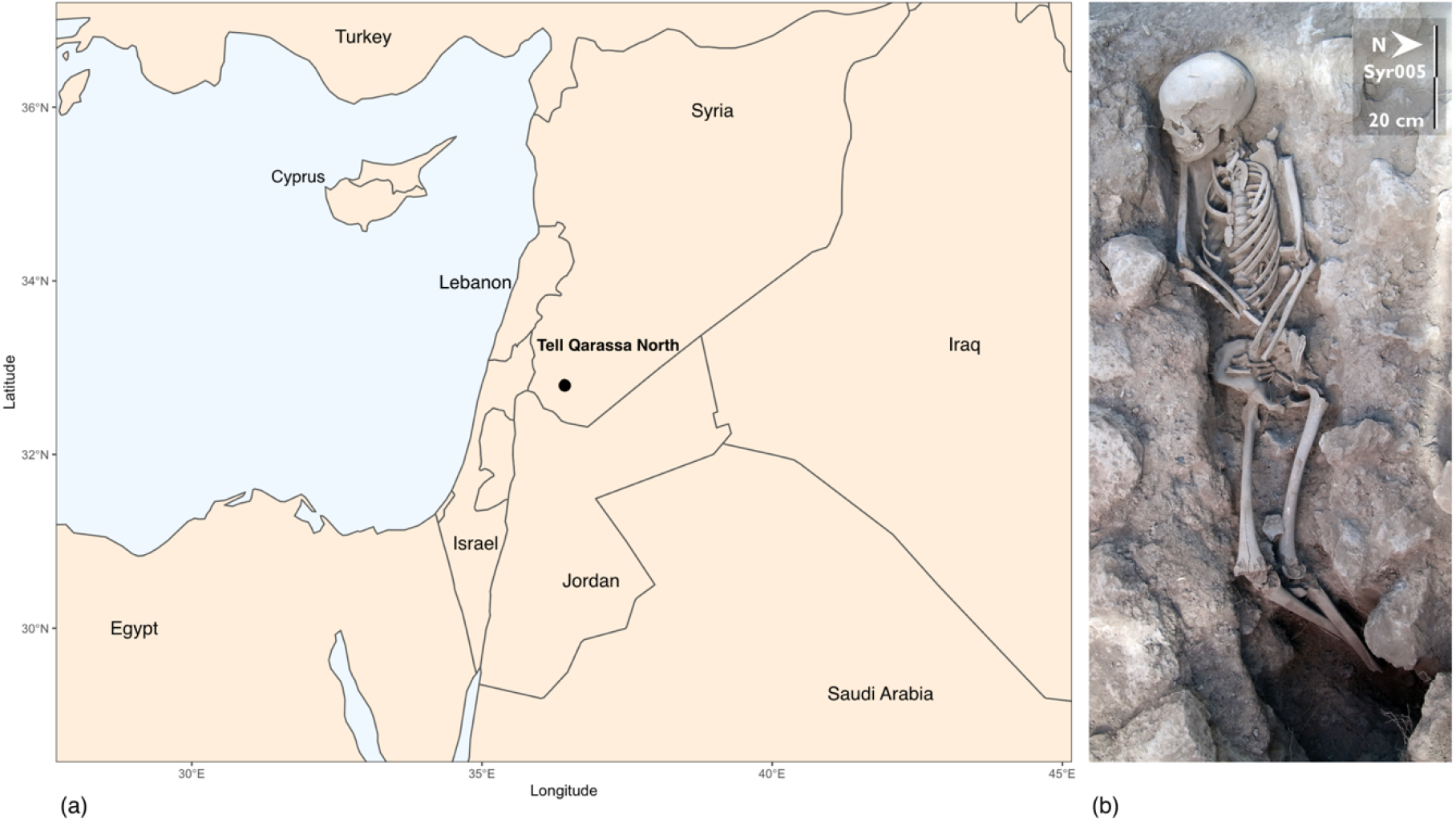
(a) Map of the Levant indicating location of Tell Qarassa in South Syria. (b) Skeletal remains of syr005 during the excavations at Tell Qarassa.

**Table 1:**
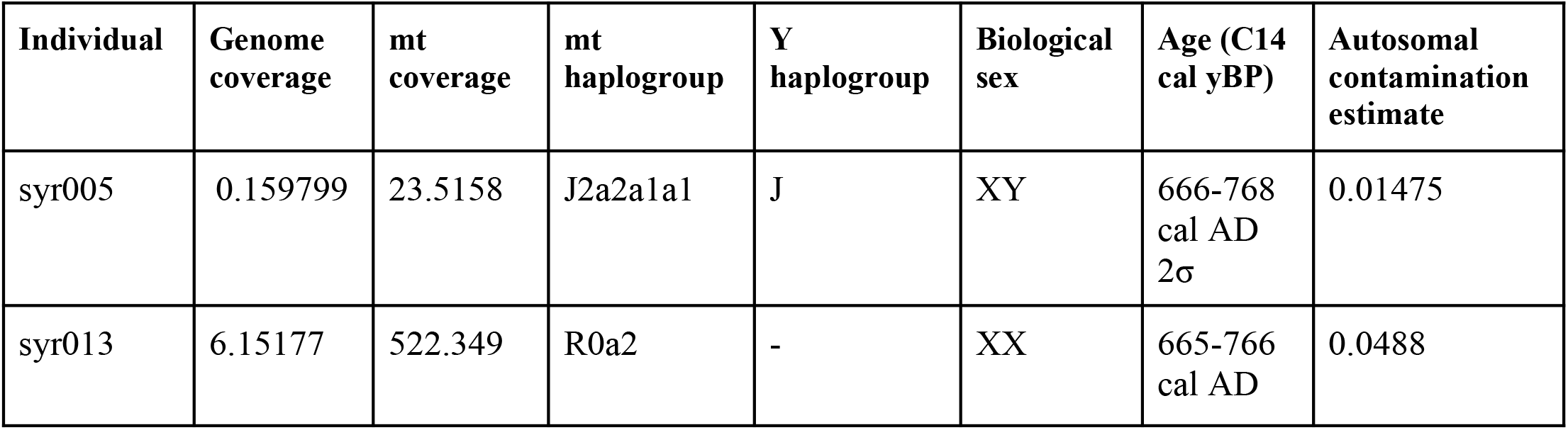
Information about the two sequenced samples.

### Genomic analysis of the individuals

To investigate the genetic identity of these two individuals, their association to past, contemporaneous and present day Middle Eastern populations as well as to shed light onto past genetic variation of Syria, a conflicted region that remains currently poorly studied, two petrous bones (syr005 and syr013) were shotgun sequenced at a depth coverage of 0.16x and 6.15x, respectively (Table 1). Sequence data from both individuals showed characteristic patterns of post-mortem damage and fragmentation, expected from endogenous ancient DNA (aDNA) molecules^22^ (Supplementary Fig. S1). We used four different methods to estimate contamination at the mitochondrial^22,23^, autosomal^24^ and X-chromosome^25^ levels and all four methods confirmed low levels of contamination (<5%, Supplementary Table S1). Two biological sex inference methods^26,27^ identified syr005 to be a male and syr013 a female. Individuals syr005 and syr013 were determined to carry mitochondrial haplogroups J2a2a1a1 and R0a2, respectively. Both haplogroups are common in the Arabian Peninsula, Near East and parts of Africa^28,29^ in concordance with the broad geographical location of the samples. In addition, the Y chromosome of syr005 was determined as haplogroup J, which is the most common haplogroup across the Middle East^30^ (Supplementary Table S2).

In order to explore general patterns of genetic affinity to modern populations, principal component analysis (PCA) was performed projecting the two newly sequenced Syrian individuals on a broad set of modern Middle Eastern, European and North African groups (Fig. 2a). The two Late Antiquity Syrian individuals fell close to Saudi, Turkish and Middle Eastern genomic variation and to some Jewish populations but do not show close genetic affinities to the geographically close Lebanese samples from^12^. Further, to obtain a better understanding of the regional variation, a second PCA was conducted, limited to 37 groups from the Middle East, Arabian Peninsula and Caucasus. Here, the Syrian samples fell close to Yemenite Jews, Saudi and Bedouins genomic variation (Fig. 2b). Notably, within these genotyped Bedouins, there are two sub-groups, both sampled in the Negev in Israel: Bedouin A and Bedouin B^2^ that were observed to have distinct distributions in the PCA: while Bedouin A seem to be more widely dispersed and overlap with other groups from surrounding regions, Bedouin B (with the exception of one individual) form a small cluster separate from all other groups in the region. From these two sub-groups, individuals syr005 and syr013 fall between the two Bedouin groups, and show a clear genetic differentiation from relevant modern-day Levantine populations (i.e Druze, Palestinian, Jordanian and Lebanese). Further, to gain insight into the genetic composition of ancient and modern populations, an unsupervised ADMIXTURE analysis was performed. ADMIXTURE was first run with a larger set of individuals (1073 individuals) from Europe and the Middle East (73 populations). For K = 2, 3 and 5, all iterations with different random seeds converged to consistent results (Supplementary Fig. S2). Therefore, we consider K=5 as a compromise between resolution and robustness of results. At K = 2, mostly north Europeans are differentiated from south Europeans, Middle Easterners and Arabian Peninsula groups. At K = 3, a new component emerged in Caucasian and Middle Eastern groups. At K = 4, another component appeared in south Europeans and Middle Eastern groups. At K = 5, a component exclusive to Middle Eastern and Arabian Peninsula groups appears (Fig. 3). This new component was seen at high proportions in Bedouins, Saudi, Yemenite Jews and our Syrian samples. Interestingly, within the Bedouins, Bedouin B was composed almost entirely of this new component. The separate cluster of Bedouin B could be the result of genetic drift, although its presence in other populations from the Arabian Peninsula suggests some degree of separate ancestry among these groups. The Late Antiquity Syrian individuals showed similar genetic composition to Bedouin B and some Saudi individuals. We conducted outgroup *f3* statistics^31^ to increase resolution on population affinities between the two Late Antiquity Syrian samples and modern Bedouins, Saudi and Yemenite Jews, as previously indicated by PCA and ADMIXTURE analysis. Both syr005 and syr013 showed the highest shared genetic drift with Bedouin B and Saudi, as expected from PCA and ADMIXTURE results. However, Yemenite Jews showed lower values, and Bedouin A were at the lower end of the spectrum among all Middle Eastern groups (Fig. 4 and Supplementary Table S3).

**Fig. 2:**
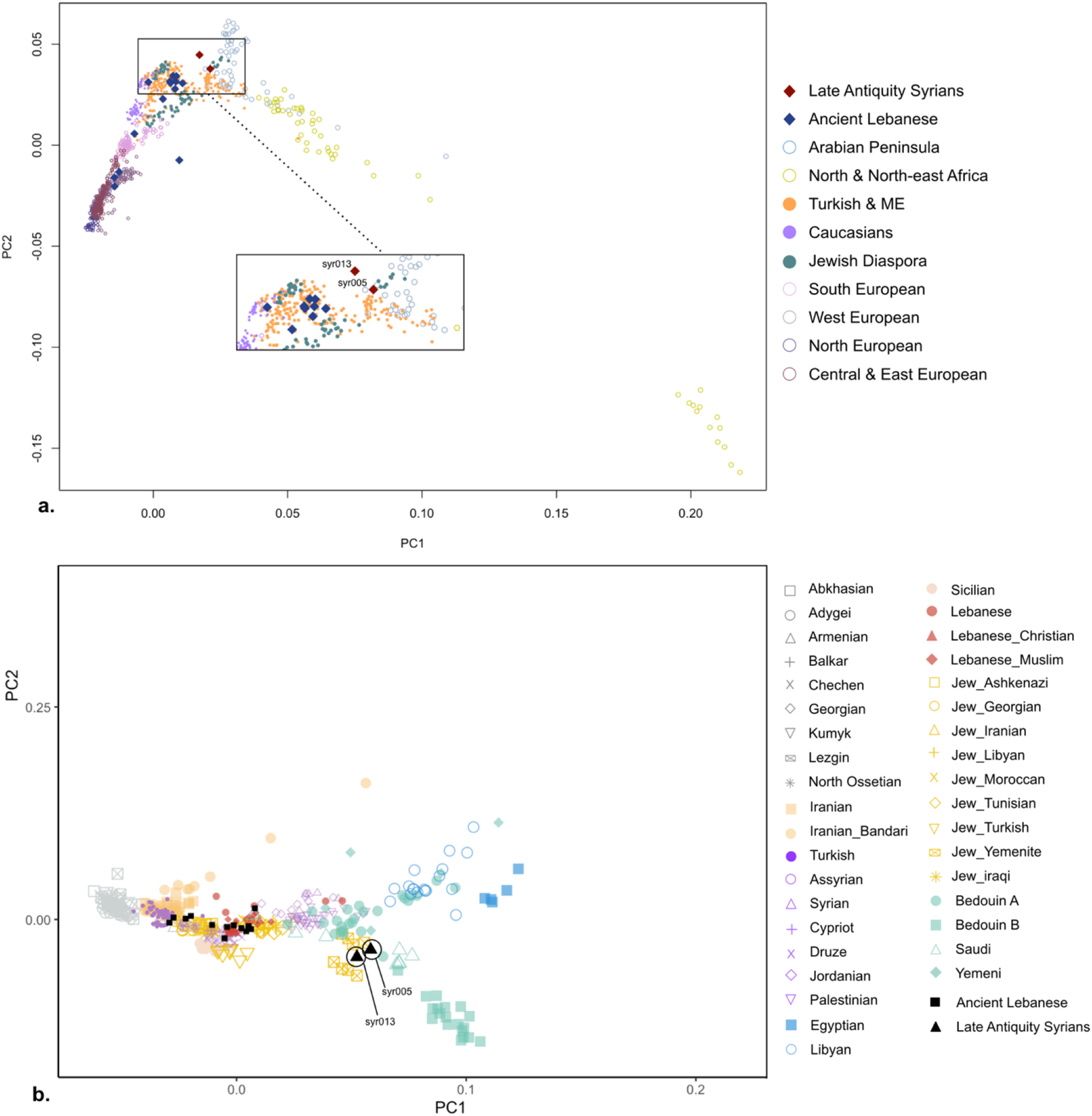
Plots showing the results of a Principal Component Analysis. Tell Qarassa individuals were projected on global variation (a) and Near Eastern variation (b). Note: One individual from Yemen falls far outside the range shown in (b). We zoom on the pattern relevant for the Late Antiquity Syrian individuals here; Ancient Lebanese include Medieval and Roman samples^12^.

**Fig. 3:**
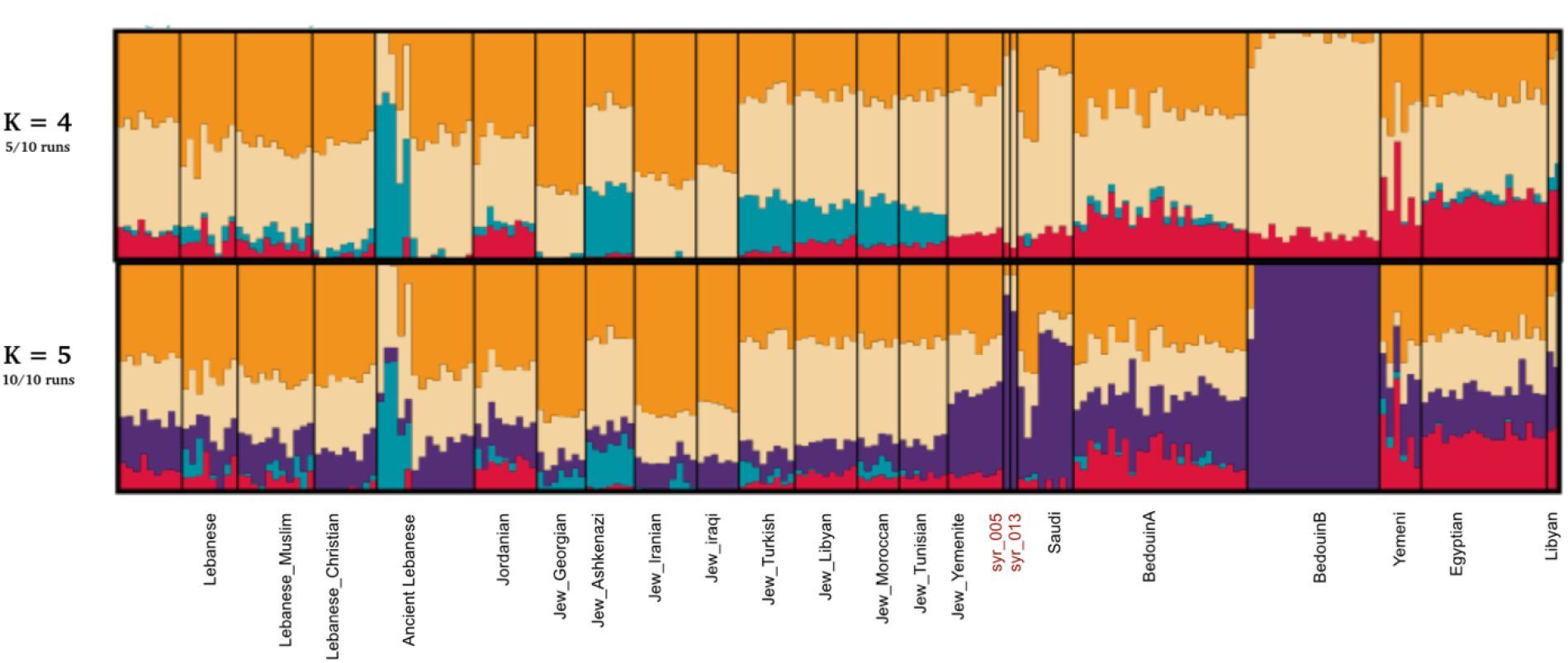
Admixture Plot at K = 4 and K = 5. K = 5 was chosen as a representative fit for the data. Late Antiquity Syrians are labelled in red (syr 005 and syr 013).

**Fig. 4:**
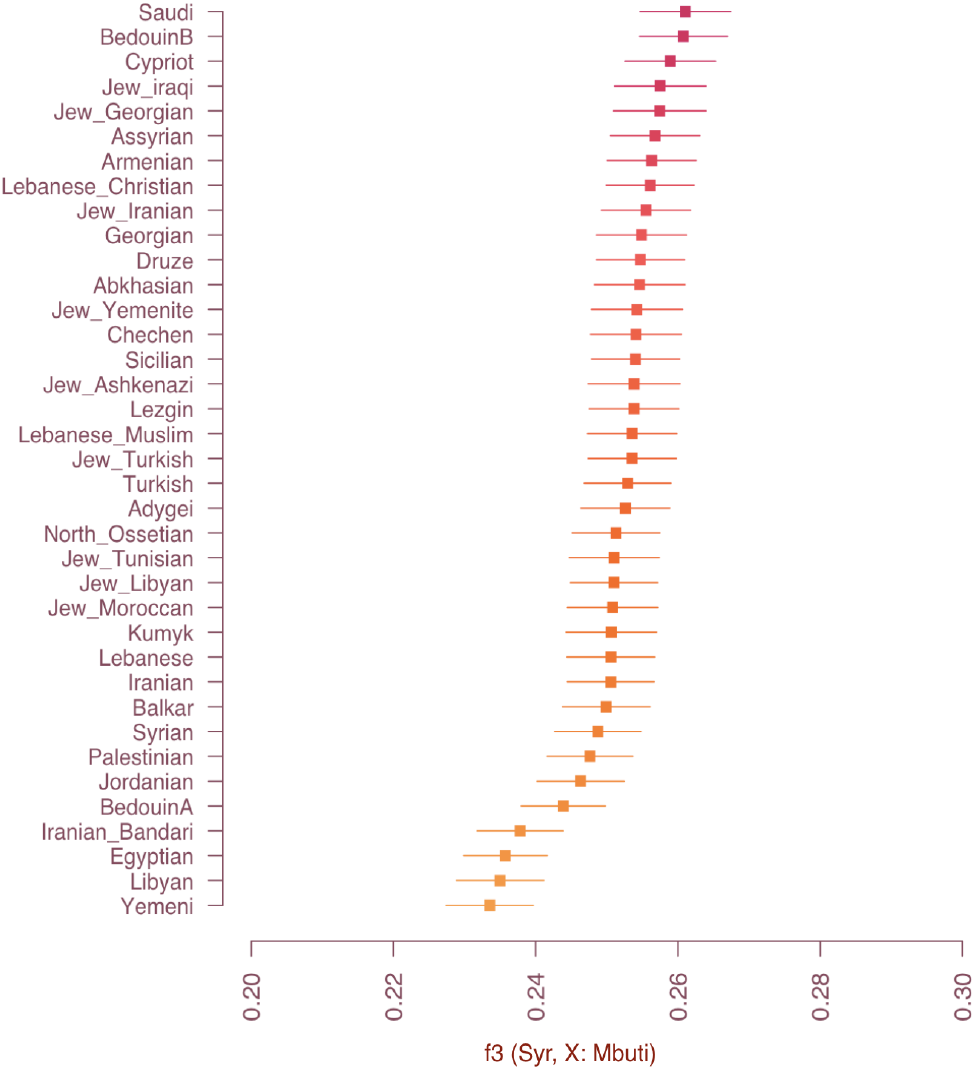
Outgroup f3 statistics showing shared genetic drift between Late Antiquity Syrians and modern-day populations.

We then tested different scenarios to investigate the most likely population affinities between our Late Antiquity Syrian samples and modern regional populations, performing D statistics of the form D (Syrian_Late_Antiquity, A; X, Mbuti) testing whether X is an outgroup with respect to the group constituted by Late Antiquity Syrian individuals and A or if there is an excess of allele sharing between A and X or the Syrians and X. When we tested Bedouin B and Saudi as candidates for groups A and X, we found that no topology was consistent with the data (Supplementary Table S4 (a-d)). Bedouins B were rejected as a sister group to the Late Antiquity Syrians to the exclusion of Saudi due to an excess of allele sharing between Saudi and the Syrians (Z=6.4) while Saudi were rejected as a sister group due to an excess of allele sharing between the Syrians and Bedouin B (Z=13.8). Testing the Late Antiquity individuals as an outgroup to Saudi and Bedouin B revealed this to be a scenario consistent with the data (Z=0.19) confirming that the two Late Antiquity Syrians form their own group and are not directly matched by any of the modern populations in our reference data.

### Estimating ancestry proportions

To obtain a deeper understanding of the different ancestries contributing to modern and historic Middle Eastern populations, we also performed a supervised ADMIXTURE analysis with sources similar to^12^. These sources represent populations close to the Eurasian and African groups contributing to modern Middle East populations. The two historic Syrian samples derive the majority of their ancestry from Neolithic Levantine populations (Levant_N, 91.3%) as well as minor proportions from Neolithic Iran (Iran_N, 4.1%) and European hunter-gatherers (WHG, 4.6%). Bedouin B are almost entirely derived from Levant_N (98.7%), with a very small percentage from Iran_N. Saudi derived more than half of their ancestry from Levant_N (63.4%), and the rest from Iran_N (36.1%) and Yoruba (0.5%). These results show that all three groups have high contributions from Neolithic Levantine groups but at different proportions, supporting the general observation of similarities between them but also consistent with a lack of a direct match to the Late Antiquity Syrian individuals in our modern reference panel (Supplementary Fig. S3). In order to gain insight into the genetic diversity of the Late Antiquity Syrians, we estimated Conditional Nucleotide diversity (CND)^32^ to be 0.205±0.003. When compared to all modern Middle East populations in our reference panel^1^, syr005 and syr013 showed the lowest levels of genetic diversity, indicating either high levels of consanguinity, small population size, close relationship between the individuals or a combination of these which would be consistent with them living in a tribal structure or being part of a small religious or cultural group. However, it should be noted that comparisons between ancient and modern populations, sequence and SNP chip data might be subject to some biases^32–34^. For comparison purposes, we included Onge (native hunter-gatherers from the Andaman Islands known to have extremely low genetic diversity due to their small population size and prolonged isolation^35^ and results showed a lower value of CND than syr005 and syr013 (Supplementary Fig. S4 and Supplementary Table S5).

### Phenotypic analysis

Finally, we analysed phenotypic traits to investigate potential lactase persistence in the newly sequenced samples (Supplementary Table S6). The SNPs analysed here are all located upstream of the gene *LCT* (encoding lactase) in introns of the gene *MCM6*, which serves as an enhancer for *LCT* transcription: −13910C/T (rs4988235), −13915T/G (rs41380347), −14010G/C (rs145946881), −13907C/G (rs41525747) and the rare variant G/A*14107 (rs182549). The variant −13915T/G (rs41380347) is common in Arabian populations, where the ancestral allele is A and derived alleles are G and C. Lactase persistence is known to be an autosomal dominant trait, thus presence of a single derived allele is sufficient for milk digestion. Interestingly, nine reads in sample syr013 were found mapping to the SNP rs41380347, out of which five were the derived allele (C) and four were ancestral (A) which indicates that syr013 was heterozygous and lactose tolerant. This could not be tested for sample syr005 as no reads covered the sites of interest. We also tested other autosomal dominant or recessive conditions reported to occur frequently in Arabs^36^ (including familial hypercholesterolemia, glucose-6-phosphate dehydrogenase deficiency, sickle-cell anemia, Bardet-Biedl syndrome etc.) but no pathogenic alleles were found in syr013 (Supplementary Table S7).

### Stable isotope analysis

The collagen extracted from the individuals was of good quality for radiocarbon and stable isotope analysis (see Supplementary table S8). The stable isotope data for these individuals are typical of a C3 terrestrial diet (δ^13^C −18.5 and −19.2; δ^15^N 13.1 and 11.5) with high animal protein intake. Consumption of food from an arid location or where manuring was practiced are also possible explanations/contributions for the high nitrogen values, but without faunal data from the period and location for comparison it is difficult to ascertain the extent of these effects.

## Discussion

The continuous development and improvement in ancient DNA methodologies and molecular techniques is constantly pushing the temporal and geographic limits of aDNA recovery, exploring older time periods^37–39^ and hostile environments (e.g. hot and humid) for DNA preservation. The Middle East and the Arabian Peninsula are pivotal regions in the timeline of human history and an increasing number of aDNA studies have attempted to understand the genetic history in these regions. Although there have been successful studies^1,3–14^, given the poor conditions for DNA preservation, this process is proving to be slower than in more environmentally favourable regions of the world. Nonetheless, given the historical importance of this region, each newly recovered DNA sequence adds an important piece to the genomic and cultural puzzle of this territory. For the two Late Antiquity individuals buried on top of a prehistoric site, we have been able to infer their genetic ancestry providing insight into the archaeological, historical and genetic record of a war-stricken region currently under difficult access and therefore heavily under-represented in recent scientific investigations. From the genetic information recovered from past individuals and that available to us today from modern populations, we can reveal that these two individuals are genetically very similar to a subgroup of modern day Bedouins (“Bedouin B’’) in the Human Origins 2.0 dataset from the Negev desert in Israel, but not to other Levantine populations such as Druze, Palestinians, Jordanians or Lebanese. In addition, they also present genetic similarities with populations from the Arabian Peninsula (Saudi and Yemenite Jews). Interestingly, when modelling the ancestry of these groups partly in the context of prehistoric groups, it becomes evident that these populations show high proportions of Neolithic Levantine ancestry suggesting long-term genetic continuity despite the dynamic population history of the region. Further, the clear presence of European (4.6%) and Neolithic Iranian ancestry (4.1%) suggests that the ancestors of syr005 and syr013 acquired these ancestries from groups in the Levant. This is further supported by the somewhat similar results seen for prehistoric individuals such as Levant_BA that also show a majority of Neolithic ancestry with equal minority proportions from Europe and Neolithic Iran.

While we cannot unequivocally identify these Syrian individuals as Bedouins, given their incomplete match and limited populations publicly available (i.e. Human Origins dataset), their high (albeit incomplete) genetic similarity to modern day Bedouin B prompts a brief discussion about the identity of Bedouins, as well as historical information known about them. Bedouins are a group of nomadic people in the Middle East historically known to have inhabited desert areas in the Arabian Peninsula, Syria, North Africa, the Sinai Peninsula, Sahara, and surrounding regions. Their traditional livelihood involves herding goats, sheeps and camels, although many groups have adopted an agricultural/urban lifestyle in recent decades. The Arabian Peninsula is described as their original home, from where they spread out north in search of pasture due to repeated droughts, growing population and tribal conflicts^40^. They have been described as being descended from two major groups, one inhabiting the mountainous regions of Yemen in southwestern Arabia and the other in North-Central Arabia^40^. They have also been labelled as ‘autochthonous’ Arabs^41^ and suggested to represent indigenous ancestral groups of the Arabian Peninsula. Despite this, they are often marginalized in practice and as such may often do not qualify for burial in one of the nearby settlements. The Leja region, a neighbouring area of Tell Qarassa, has a long history of occupation by Bedouin nomads and it is known that the area of Tell Qarassa was occupied by Bedouin of the Banu Sarma in the sixteenth century^42^. Because of the long-term continuity of the Bedouin nomadic lifestyle, burial practices of today or the recent past are regarded as comparable with the distant past back to the early years of Islam^43^ (Supplementary Text S1 – Bedouin Burials).

Given their practice of consanguinity and strong barriers to extra-tribal marriage, they have low genetic diversity, a small effective population size and a high incidence of recessive disorders, which has made them subjects of clinical studies in the region^36,44^. The fact that the genotyped Levantine Bedouins available in the Human Origins dataset all stem from the Negev desert in Israel but show highly different genetic ancestry (Bedouin A versus Bedouin B)^2^ highlights that there might be a substantial degree of genetic structure within Bedouins in the region. However, the full extent of this structure might not be fully understood due to a lack of available past and present genetic data. Studies on modern Bedouins elsewhere, e.g. in Qatar report them as ‘coldspots’ of admixture in the Peninsula compared to other populations^45^. It is suggested that the Negev Bedouins originate from a small founder population and most of their ancestors migrated from the Arabian Peninsula to the Negev and Sinai regions around 700 CE, i.e., shortly after the spread of Islam^46^. Similar migration events have been recorded, including for the Tell Qarassa region, which was controlled by the city of Bostra, an important settlement in southern Syria during the Roman and Byzantine empires^47^. This area was occupied by the Ghassanids, a nomadic group from the South Arabian Peninsula that arrived at the end of the fifth century and became Byzantium’s principal Arab ally^48^ during the early sixth century, ruling a large Christian population. This period saw an influx of Islamic Arabs along with other migrants, many of which maintained a nomadic lifestyle^49^. It is clear that the Late Antiquity in the Levant, were highly dynamic times driven by Arab migrations to the Levant, mirrored by clan-structured groups such as Bedouins. Within this framework, our two Syrian samples could potentially represent part of one of these highly structured nomadic tribes that arrived in the Levant from Arabia under strict cultural, social and religious rules.

The Tell Qarassa graves differ from most other excavated examples of early Islamic burials because they were not located in a cemetery and – from the available evidence-do not seem to have been located near a permanent settlement of the period. From studies of nomadic Bedouin it appears that when someone dies they are buried immediately in a prominent nearby location^50^. There are a number of aspects of the Tell Qarassa burials which can be documented in Bedouin burials both from the recent and more distant past (Supplementary Text S1 – Bedouin Burials). Based on this information and also taking into account the genetic results it seems likely that the deceased were Bedouin.

The collagen stable isotope data from these two Late Antique individuals are difficult to interpret as there are no contemporary data from this region with which to make a direct comparison. Data from Syria (Northern) and nearby Northern Jordan from periods before 651AD (Late Roman/Byzantine, Parthian) and some modern samples, demonstrate similar δ^13^C values, but generally lower δ^15^N (7.3-9.0) than the two Late Antique individuals studied here (see Supplementary table S8 and references therein). Dietary stable isotope studies of ancient Islamic individuals have been widely studied in Iberia (and the Balearic Islands), but less extensively elsewhere. Lopez-Costas and Alexander (2019)^51^ provide a useful review and discuss that there is no strong evidence that Islamic diet has a distinct isotopic signal that distinguishes it from contemporary Christian diets (e.g. Dury et al., 2019^52^), despite some well-known cultural differences (e.g. the prohibition of pork in Islam, fasting and fish eating in Christianity). As Lopez-Costas and Alexander (2019)^51^ point out further research is required to investigate such distinctions and certainly further work is required in the Middle East to expand our knowledge of dietary stable isotopes in the Late Antique period.

### Genomic ancestry of Levantine populations

From a genetic perspective, Levantine populations today fall into two major groups: one sharing genetic similarity with modern-day Europeans and Central Asians (Eurasian cline), and the other closer to Middle Easterners and Africans (African cline)^13,14^. Generally, the ancestry proportion contributed by Neolithic Levant populations increases towards the Middle East and Arabia, while there is more Neolithic Iran ancestry in groups in the Levant and the Caucasus. Western Hunter Gatherer (WHG) proportions increase towards the north-west (i.e., towards Europe), and also in populations located on the Mediterranean coast. In the Arabian Peninsula and North Africa, the proportion of African ancestry increases. Different archaeogenetic studies have highlighted the complex demographic changes that have shaped genomic ancestry in the Levant^1,3–14^.The majority of prehistoric and historic genomes sequenced so far follow a similar pattern, but several studies have also found single individuals deviating from this pattern^6,8,11^. Such finds suggest additional small-scale migrations and the presence of structure within the region that is not displayed in the majority of the past society. Our Late Antiquity Syrians and Bedouin B are such exceptions to the general genetic pattern as they show an extremely high Neolithic Levant ancestry, which is higher than most neighbouring Middle Eastern groups (Fig. 5). The high level of Neolithic Levant ancestry observed in syr005, syr013 and Bedouin B denotes a certain level of continuity of this component, at least in a few groups of this region.

**Fig. 5.**
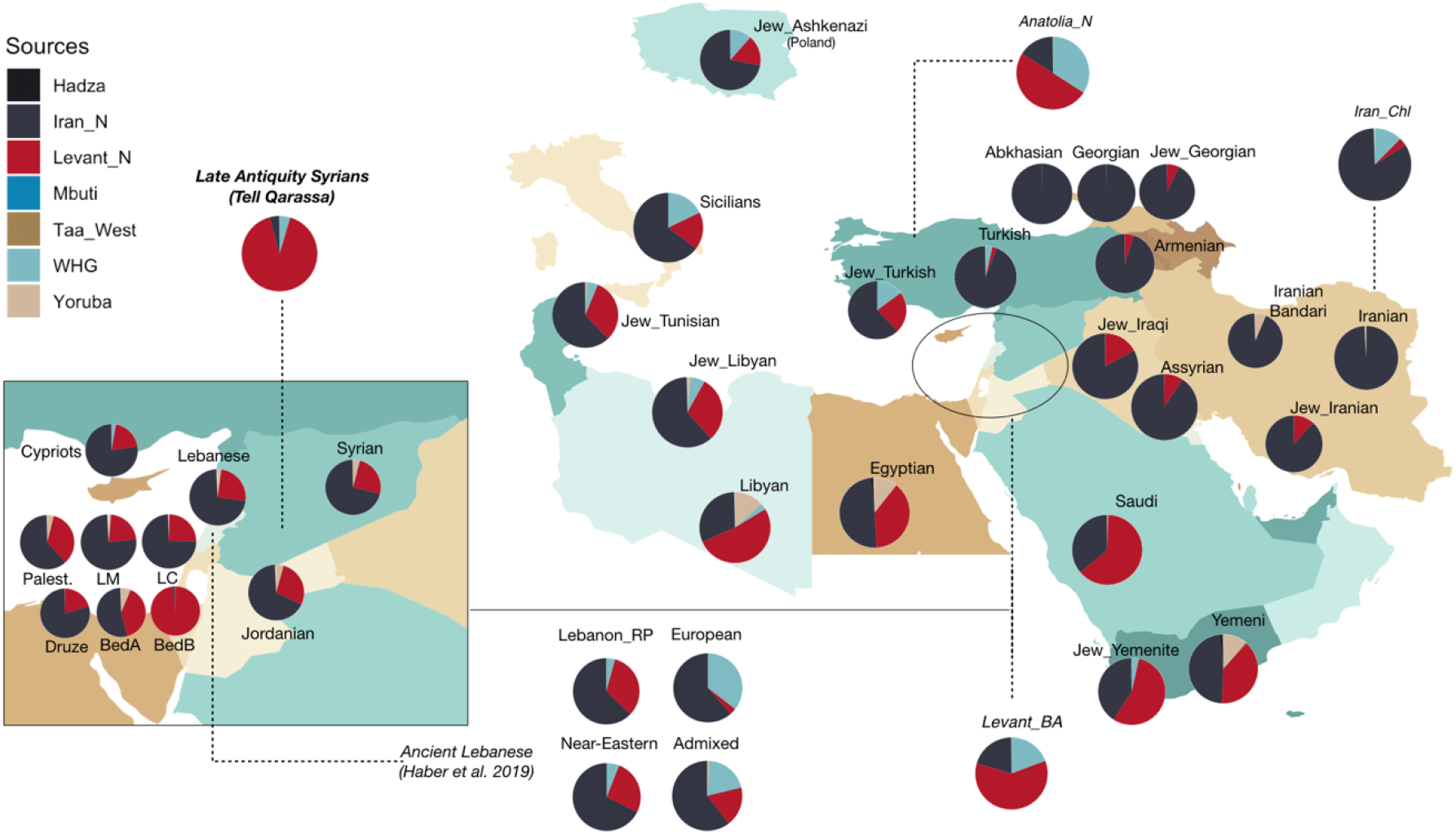
Supervised admixture proportions projected on regions of sampling. Ancient populations are joined by dotted lines to the historical region of origin, with Late Antiquity Syrians highlighted in the top left. Levantine populations have been zoomed in for clarity (Palest.: Palestinians, LM: Lebanese Muslim, LC: Lebanese Christians, BedA,B: Bedouin A, B). Ancient Lebanese individuals from^12^ are also shown, with labels corresponding to those used/suggested in the study.

The small genetic differences observed between these two groups suggest a slightly different trajectory and contacts with other groups since the Neolithic. The clear distinction between syr005 and syr013 from modern Levantine populations combined with the evident genetic similarity to Neolithic Levantine individuals posits an interesting genetic picture that can be explained under two alternative scenarios: 1) a long-term continuity followed by genetic isolation due to strong social, cultural and religious barriers resulting from the Islamisation of the region during the Early Islamic period or 2) the result of a past migration from the Arabian Peninsula. Under the first scenario, syr005 and syr013 would exemplify a group representing long-term local continuity since the Neolithic that would have converted to Islam relatively early after the Arab expansion took place. Hence, the original genetic structure of the region would have been conserved due to strong social, cultural and religious barriers over time preventing mixing with neighbouring populations, resulting in the highly drifted population observed in our data. Long-term genetic continuity has been suggested for various Levantine populations, such as for modern Lebanese which are found to be genetically similar to Bronze Age populations from the region^9^, as well as for present-day populations which have been reported to retain ancestry from local populations despite admixture events such as the Crusades in the Middle Ages^12^. Under the second scenario, our data is best explained by migration from the Arabian Peninsula to the Levant, a scenario concordant with historical records of a migration from the Arabian Peninsula to the region not long before the time of our two Late Antiquity Syrians (i.e. 7^th^ and 8^th^ centuries) (see above and Supplementary Text S1 – Early Islamic Southern Syria). Moreover, the presence of Neolithic Levantine ancestry in modern-day populations from the Arabian Peninsula seems to support this scenario. If we assume that Neolithic Levantine ancestry was already present in the Arabian Peninsula during historic times or earlier, we would expect to see a reintroduction of Neolithic Ancestry into the Levant where this would have been diluted due to the multi-layered history during the previous millennia. The remarkable affinity to Arabian and Neolithic ancestry would have been maintained overtime as a result of the traditional clan structure of Arab tribes.

In a region that has a complex history of population movement and mixing, the lack of admixture presented by these samples is surprising. One explanation for this could be that these individuals come from an endogamous society, i.e. one that culturally restricts sexual relations to within the society itself or similar groups. Such endogamous practice would likely be consistent with the social practices of the expanding Muslim population of this time. Moreover, the burials of these individuals have traits of Muslim burials. It is possible that we have found some of the earliest evidence for an endogamous Muslim population in the region.

Our conclusions are limited by the availability of genomic data both from modern and prehistoric populations. The fact that one can observe substantial genomic structure among the few genotyped Bedouin individuals^2,46^ suggests that there is a significant level of hidden genetic structure in sampled and unsampled groups. Extensive additional sampling from these groups is crucial to our understanding of the extent of their genetic structure today, and potentially in identifying relatively genetically isolated populations, which has implications for population genetic and clinical studies. The Middle East is a region with a complex history and a diverse ethnic and genetic composition, and our current understanding of the genetic structure in the past and present appears to have only scratched the surface.

### Conclusions

Genomic methods are a powerful tool to study an individual’s ancestry which informs us about past demography and population dynamics, but not directly about the cultural identity of the individual in the absence of archaeological context. The genetic results of our Late Antiquity individuals uncover higher genetic affinities to some modern Bedouin groups, considered representatives of ancestral Arabian indigenous groups, rather than to other nearby Levantine populations.

We complemented the genomic data with additional archaeological and historical information to obtain a clearer overview of the two individuals from Late Antiquity Syria. The archaeological record reveals that these two individuals were deliberately buried on top of a Prehistoric burial site during the Late Antiquity period following an Islamic funerary ritual, but not within a traditional Muslim context, i.e. a traditional Muslim cemetery (see Supplementary Text S1 – Muslim Burials, Bedouin Burials and Re-use of Ancient Sites). Within this framework, it is possible that these two individuals were transient Muslims in the region. From the proximity of the ^14^C dates and the fact that these are the only non-prehistoric burials at the site, it seems plausible that they died at the same time or shortly one after the other. The close proximity of the two graves suggests that the two individuals were related, and while our genetic results do not show first degree relationship, we cannot discard another level of kinship. The absence of trauma to the bones and the young age of the deceased suggests that they may have died from a disease, possibly the Justinian plague which ravaged the Middle East from 541 AD to 749 AD recurring in cycles^53^. Specifically, the dates of the burials may be linked to the outbreak of 79 AH (698 AD) which was reported in Syria by as-Suyufī^54^.

The genomic ancestry of the two people buried at Tell Qarassa in the late seventh or early eighth century offers a tantalising glimpse of early Islamic society in Syria. This study provides a further insight into the re-use of prehistoric sites as cemeteries by Muslims and in particular Bedouin. On a general level, this burial provides additional evidence for the early adoption of specific Islamic burial rites which were followed even in remote locations. At present there are no examples of genetic studies from the region which relate to this period, the only genetic data which relates to early Islamic burials is the study of two individuals from the south of France^55^. Our results show early presence of Muslim Arabs in the Syrian countryside and provide more evidence of the Bedouin contribution to early Islamic society.

## Materials and Methods

### Archaeological context

The archaeological site of Tell Qarassa, located in modern-day Syria (Fig. 1a), is prominently known for its evidence of human occupation since the Epipaleolithic period to the Iron Age^56–58^. Located on the shores of an ancient dried up lake, it is described to consist of multiple sites or so-called ‘Tells’, one of which (Northern Tell) contains remains from Pre-Pottery Neolithic B to Late Chalcolithic settlements, while the other (Southern Tell) holds evidence of Early Bronze to Iron Age remains. Evidence of a Natufian settlement has also been found in close proximity^59^. The village of Qarassa is a Druze community today. Pre-Pottery Neolithic mortuary practices have been described from this site, shedding light on such practices in this period^18,19^. The Late Antiquity burials consisted of two narrow graves that were opened in superficial layers of Tell Qarassa North. While in the Neolithic burials the bodies are placed in flexed position, these bodies were placed in decubitus position, oriented east-west, with the head at the west, facing south. Individual syr005 was a 14-15 years old male (Fig 1b) and individual syr013 was a female of about 15-21 years at the time of death (Table 1). Age-at-death estimation was based on the following criteria: pattern of dental eruption, synostosis of epiphyses on the long bone and closure of the sternal ends of the clavicles^60–62^. These individuals were located very close to each other. However, intensive archaeological fieldwork has not yielded evidence of further burials from this period at the site. Therefore, the archaeological record does not support that these burials belong to a cemetery for a specific community. It is suggested that these burials may belong to nomadic populations, pilgrims, or plague victims^21^.

### Radiocarbon dating and Isotopic analysis

Collagen was extracted from two petrous bones following a modified Longin method^63,64^ at the Molecular Archaeology Laboratory at La Trobe University using cold 0.6M HCl and yields are recorded in Supplementary table S8. The collagen was directly radiocarbon dated (AMS) at Waikato University in New Zealand and calibrated using the Oxcal 4.3 program^65^, and the IntCal13 calibration curve^66^. Radiocarbon results for individuals syr005 and syr013 were 666-768 Cal AD 2σ (1294 ± 18 Cal BP, Lab code wk-46474), and 665-766 Cal AD 2σ (1302 ± 15 Cal BP, Lab code wk-46475), respectively. The collagen stable isotope values and quality parameters were also measured at Waikato and are presented in Supplementary table S8.

### Sample Preparation and Sequencing

Prior to DNA extraction, the petrous bones were UV irradiated (6 J/cm2 at 254 nm) and the first millimetre of bone surface abraded using a Dremel™ tool. DNA was extracted from a 100-200 mg piece of the bone using a silica binding method^67^, with an incubation of 24-48 h, using the MinElute column Zymo extender assembly replaced by the High Pure Extender Assembly (Roche High Pure Viral Nucleic Acid Large Vol. Kit) and performed twice for each sample. Further, blunt-end Illumina multiplex sequencing libraries were prepared^68^ resulting in two double stranded libraries per sample. Library amplifications were performed as in^69^ using indexed primers^68^ and 4 −11 cycles (11 and 10 cycles for syr005; 4 and 6 cycles for syr013). All extraction and library preparation steps were conducted at the dedicated ancient DNA facility at Stockholm University. A total of four DNA libraries were shotgun sequenced on a HiSeq X10 sequencing platform (150 bp paired-end reads) at the NGI Stockholm.

### NGS Data Processing

Sequenced reads were mapped to the human reference genome build 37 (hs37d5) using BWA aln^70^ with non-default parameters −l 16500 −n 0.01 −o 2. Data was merged at library level using samtools v1.5^71^. PCR duplicates were collapsed using a modified version of FilterUniqSAMCons_cc.py^33,72^. Different libraries for one individual were then merged into one bam file using samtools v1.5^71^. Reads shorter than 35 bp, showing more than 10% mismatch with the reference and/or a mapping quality score below 30 were discarded. Biological sex was inferred using two different approaches^26,27^.

### Contamination estimates

Contamination estimates for the two samples were assessed on three different levels: mitochondria, X-chromosome and autosomes. Two methods were used to estimate mitochondrial contamination^22,23^. X-chromosomal contamination was estimated for the male individual syr005 using the approach implemented in ANGSD^25^. VerifyBAMID^24^ was used to estimate autosomal contamination in both samples. It uses a hypothetical ‘true genotype’ model and checks whether reads in a .bam file are more likely to match a single individual or result from a mixture of other samples, such as a closely related/a different individual.

### Uniparental haplogroups

Mitochondrial haplogroup identification was carried out using Haplofind^73^ and HaploGrep^74^, two web-based applications that use the Human Phylogenetic Tree^75^ To identify the Y-haplotype of sample syr005, reads mapping to the Y chromosome with minimum base and mapping quality of 30 were compared to biallelic substitution SNPs from the International Society of Genetic Genealogy (ISOGG, https://isogg.org/). We excluded A/T and G/C SNPs to avoid strand misidentification and C/T and A/G SNPs to avoid post-mortem damages.

### Population Genetic Analyses

For population genetic analyses, we merged the ancient samples with modern genotype data of the Human Origins 2.0 dataset^1^. We generated pseudo-haploid representations of the ancient individuals by randomly drawing one allele from the samtools mpileup^71^ output at each SNP site, restricting the analysis to minimum mapping and read qualities of 30 and coded transition sites as missing to avoid post-mortem damage.

A principal component analysis was conducted using smartpca^76^ using the options lsqproj and shrinkmode. The first PCA included populations from North, West, South, Central and East Europe, Caucasus, Turkey and the Middle East, North and North-east Africa, the Arabian Peninsula, Medieval samples from^12^ and the ancient Syrian samples from this study. In order to identify the closest modern populations to the ancient Syrian samples, we generated a second PCA with increased resolution by reducing the original set of modern populations. For this second PCA we excluded divergent groups such as Europeans, Somalis and was limited to 37 groups in the Middle East, Turkey and Caucasus (493 individuals, henceforth Middle East (Middle Eastern) panel). PCA plots were generated using GNU R.

We used ADMIXTURE^77^ for a model-based clustering analysis. All data were haploidized by randomly picking one allele per individual at each SNP. Next, the dataset was thinned by pruning out SNPs in linkage disequilibrium using PLINK v1.9^78^ with a window size of 200 kb, a step-size of 25 and a squared correlation (r^2^) threshold of 0.4. Following this step, 87098 of 602366 variants were removed. ADMIXTURE was run for 10 iterations with different random starting seeds, five-fold cross validation and the number of ancestral populations (K) was varied from 2 to 10. Results of admixture were visualized using Pong^79^. We first ran unsupervised analyses to study assignment of genetic clusters in the larger set of 73 populations from Europe, Middle East, Arabian Peninsula, North Africa and Caucasus (1073 individuals). Further, we ran supervised ADMIXTURE using another set of populations that included ancient groups from Europe, Middle East and a few additional African groups to quantify ancestry proportions of our samples in a more historical context. A new dataset was created by merging SNPs from various ancient populations that represented more historical periods in the Levant and adjoining areas as well as more representative populations from Africa: Western European Hunter Gatherers (WHG), Neolithic populations from Anatolia, Levant, Iran (Anatolia_N, Levant_N, Iran_N), Bronze Age (Anatolia_BA, Levant_BA, Arm_MLBA), Iran_Chl, Arm_Chl (Arm: Armenian; MLBA: Middle-late Bronze Age; Chl: Chalcolithic), Natufians^1,2,80–83^ as well as some modern African populations (HumanOrigins 2.0)^1^. This choice of source populations was initially based on those used by^12^ but we reduced the source populations to WHG, Levant_N, Iran_N, Yoruba, Taa_West, Mbuti and Hadza as our main focus is on the ancestry in two Levantine individuals. ADMIXTURE was run in the supervised mode using these seven populations as references (K = 7) with 10 iterations, random starting seeds and five-fold cross validation. Finally, an unsupervised admixture (K = 2 to K = 10) of the same dataset was also run to compare the results of both analyses (see Supplementary Text S2 and Supplementary Fig. S5).

To detect shared genetic history between our samples and various Middle East populations we used *f*_3_ and *D* statistics^31,84^. Population genetic summary statistics were calculated using POPSTATS^85^ with sub-Saharan Mbuti rainforest hunter-gatherers as an outgroup. Standard errors were estimated using a weighted block-jackknife procedure.

To estimate genetic diversity in our data, we used conditional nucleotide diversity (CND)^32^. CND is a measure of genetic diversity within a population based on a comparison of nucleotide differences between two individuals. We used transversion sites that are polymorphic in the Human Origins SNP array. Standard errors were estimated using a block jackknife procedure.

### Phenotypic Analysis

Given the observed Arabian ancestry in our samples, we looked at phenotypic variants that might be expected in this region. For instance, lactase persistence (LP) has been described from the Arabian Peninsula^86^. Given that Bedouins are known to be camel herders, we speculated our samples to have this LP variant. We also looked for common recessive disorders described in Arab populations from the OMIM (Online Mendelian Inheritance in Man) catalogue^87^. Information about SNP positions and the phenotype was taken from dbSNP^88^ and SNPedia^89^. A pileup file was created in samtools from .bam files with a high base quality (q > 30) as well as mapping quality (Q > 30) and the occurrence of variants known to be involved in LP and other disorders was checked.

## Acknowledgments

This work was funded by a grant from the Royal Physiographic Society of Lund (Nilsson-Ehle Endowments) to TG and CV and the La Trobe Internal Research grants to CV. MS was part of the Erasmus Mundus Master Programme in Evolutionary Biology (MEME). TG is supported by a grant from the Swedish Research Council Vetenskapsrådet (2017-05267). MJ and AG were supported by grants from the Swedish Research Council Vetenskapsrådet and the Knut and Alice Wallenberg foundation. JS is supported by a grant from Marie Skłodowska-Curie Actions (European Commission, no. GA 750460; H2020-MSCA-IF-2016). The archaeological research at Tell Qarassa was supported by the Spanish Ministry of Economy and Competitiveness (grant HAR2016-74999-P) and the Palarq Foundation.

We thank Arielle R. Munters for initial data processing, Phillip Edwards for fruitful discussion and Frank Braemer for comments on an early draft of the manuscript. The computations were performed on resources provided by SNIC through Uppsala Multidisciplinary Center for Advanced Computational Science (UPPMAX) under projects sllstore2017020 and 2019/8-150. Shotgun sequencing was performed at the National Genomics Infrastructure (NGI) Stockholm.

## Author contributions

T.G. and C.V. conceived the study. H.B. I.U. and C.V. performed sampling and lab work. M.S. and T.G. performed analyses. J.S., E.I., A.P., J.J.I., F.B., M.J. and A.G. provided either archaeological material and background, or input about genomic analyses. M.S., T.G. and C.V. wrote the manuscript with input from all co-authors.

## Additional Information

The authors declare no competing interests.

## Data availability

Generated sequence data is available at the European Nucleotide Archive (ENA) under the accession number PRJEB38008.

## Notes

### Competing Interest Statement

The authors have declared no competing interest.

