## Supplementary Information for "Bioarchaeological analysis of one of the earliest Islamic burials in the Levant"

**This supplement contains**

Supplementary Text S1
Supplementary Text S2
Supplementary Figures
Supplementary Tables
References in the Supplementary Text

### Supplementary Text S1

#### Historical Discussion of Tell Qarassa Burials

##### Muslim Burials

Muslim graves are very diverse yet easily distinguishable from other forms of religious burial. The diversity is mostly concerned with grave markings, traditions around the funeral and the precise form of the grave pit itself. The defining features of Islamic burials are the position of the body, the lack of a wooden coffin and the speed of burial. Despite the fact that much of the focus of Near Eastern Archaeology has been on funerary remains surprisingly few Islamic burial sites have been investigated. Of the excavated sites, few have been published and even fewer have been subjected to analytical tests. The rarity of archaeologically investigated Muslim burials is largely the result of religious sensitivities and traditional beliefs about the necessity for having a complete body with which to enter the afterlife (for a detailed discussion of the archaeology of Muslim burials see[1]).

Although death and burial are processes which are nearly always connected with religious belief the Quran has very little to say either about funerals or how a body should be buried (see Quran 5:31, 9:84 and 21:35). Instead almost all aspects of Muslim practice around death and burial are derived from traditions (*hadith*) and religious reasoning (*ijtihad*). *Hadith* (or traditions) are sayings or actions which are traditionally believed to have come from the Prophet Muhammad. As the example of a perfect Muslim, Muhammad's actions are necessarily in accordance with Islam. Because Muhammad himself did not write down his thoughts or pronouncements, most of the traditions originate from one or other of his close associates who transmitted them orally until such time as they were written. Later the various traditions were collated and organized into four schools (*madhab*) based on their original source and form of transmission. The actual difference between schools is quite small and collectively the different traditions form a body or religious knowledge known as the *Sunnah* (compilation). However, the hadith themselves give limited information about funerary rites and what to do with the body of the deceased and for detailed guidance, Muslim scholars have turned to religious reasoning (*ijtihad*) from which they derive rules based on both the Quran and the traditions[2]. During the thirteenth and fourteenth centuries much of this information was collected in funerary manuals which give specific information on how a dead person should be prepared for funeral and buried[3].

The basic requirements of a Muslim burial as described in Islamic texts are:

1) the deceased should be buried before nightfall on the day of death and if the death occurs too late in the day or at night the burial should take place before sunset the next day.

2) the body should be washed prior to burial and this should usually be carried out by close relatives of the same sex,

3) there is no need for a coffin (although corpses are sometimes carried to the grave within a coffin which is then retrieved for re-use) and the dead should be buried either wrapped in a shroud or where circumstances do not permit in their own clothes.

4) the body should be laid in the grave, usually on their right side, with their head turned to face Mecca. The body was aligned perpendicular to the *Qibla* axis so that only the head faces Mecca. In practical terms this means that in Syria and the Levant (Jordan, Lebanon, Palestine/Israel) the body will be aligned East-West with their face turned towards the south.

5) one of the defining features of Muslim burials is the rule of only one person per grave. This means that husbands and wives are not buried together, and also collective family tombs are forbidden. Very occasionally and in extreme circumstances this can be relaxed for victims of plague or warfare.

According to Muslim tradition, burials and cemeteries should be located outside the inhabited areas usually on the outskirts of a town or village. Visitation of graves by relatives or others is discouraged particularly in fundamentalist Sunni Islam although it is widely practiced and in some cases is an established tradition.

The shape of the burial pit varies, although two main types were commonly used – a single rectangular trench dug out of the ground with a smaller human sized trench at the bottom (*shiqq or shaqq*) or a rectangular trench with a niche in one side where the body is deposited (*lahd*). Another grave form used occasionally is the shaft tomb comprising an underground chamber linked by a tunnel either directly to the surface or via a shallow pit. It seems likely that the different forms of burial originated either at different times and/or in different places.

There is currently little information on this and the small number of archaeologically excavated Muslim burials is too small to draw any conclusions. However, from the very few excavations of early Islamic (pre-ninth century) graves carried out so far, it is evident that simple rectangular burials were the earliest form used with examples uncovered in France[4], Spain[5] and Palestine/Israel[6,7]. Shaft tombs have also been attested from the early eighth century at the site of Nahal ‘Oded in the Negev[8]. The earliest archaeologically investigated example of a grave with a side niche (lahad) is from Çatal Höyük in central Anatolia and is dated to the Seljuk period (approx. 11th-13th centuries) or later[9,10].

**Archaeology of Muslim Burials**

Whilst the number of scientifically excavated Muslim graves is small, the number of early Islamic graves is even less. For example, archaeologists excavating the early Islamic palace at Qastal in Jordan discovered a graveyard dating to the early Umayyad period (late seventh or early eighth century AD) which is of considerable significance for understanding early Islamic society. However, religious sensitivities meant that it was not excavated[11]. In most cases the anthropological investigation of early Islamic human remains is only carried out when they are found unexpectedly such as the individuals excavated at Qasr Hallabat who appear to have been murdered[12]. In non-Muslim majority countries, there is usually less reluctance to excavate graves thus the largest number of archaeologically documented Muslim graves comes from the Iberian Peninsula and to a lesser extent from Israel. In Spain and Portugal where the Muslim presence ended more than 500 years ago, a large number of Islamic cemeteries have been excavated. In addition, a wide range of analytical methods have been employed including isotope analysis to determine diet and geographical origins[5], measurement of physical characteristics to investigate ethnic identity[13], and taphonomic investigations to understand post-depositional preservation or remains[14]. In Israel, excavations at the early Islamic city of Ramla uncovered many early Islamic graves including a large cemetery that was subsequently built-over in the ninth century[6,7,15]. The sandy nature of the soil in Ramla meant that the sides of the rectangular graves were lined with stone walls[7]. There has been some anthropological investigation of the skeletons discovered in Ramla although this is rare[16]. Political sensitivities about human remains mean that for most early Islamic burials discovered in Israel, palaeo-anthropological studies have been limited to recording the position of bodies and the presence or absence of bones.

**Early Islamic Southern Syria**

Although there had been previous Arab incursions into Syria, the final Muslim conquest of Syria took place in 630's with the capture of Bosra in 634 and Damascus in 636[17]. In 661, Muawiyya, the governor of Syria succeeded to the caliphate and became the first of a series of Umayyad caliphs to rule the Islamic world from Syria. During this period, Damascus was established as the capital and cities throughout Syria and Palestine were developed as centres of Umayyad rule. Also, during this time, the Umayyads and their clients built a series of palaces along the desert fringes of Syria and Jordan[18]. Bosra was the de-facto capital of southern Syria and in the ninth century Ya'qubi described it as the capital of the Hauran as an important area for wheat cultivation[19]. Other important cities in the region included Suwayda, Ezra, Dera and Shehaba.

Tell Qarassa lies at the southern edge of the *Leja,* a forbidding basalt covered region located between the ancient cities of Damascus and Bosra. Although the village of Qarassa appears to be a relatively recent settlement there is evidence of Roman, Byzantine and Medieval occupation both in the immediate area and in the wider region. Archaeological survey work in the region indicate that there was substantial population during the Roman, Byzantine and early Islamic periods with a significant break at the end of the Umayyad period (circa 750). Large scale occupation of the area appears to have resumed in the eleventh and twelfth centuries and continued into the early Ottoman period[20]. Sixteenth century Ottoman tax registers indicate that the Leja region was occupied both by villages and nomadic groups including Turcoman and various Arab Bedouin tribes[21]. The first Druze populations arrived in the region in the seventeenth century fleeing persecution in Mount Lebanon.

The closest historical settlements to Tell Qarassa are Busra al-Harir (6km to the west), Najran (3.5km to the east) and Harran (6km to the north) each of which has significant Roman, medieval and later remains. In addition, there is the small settlement of Duwayra which although not ancient was already in existence in the sixteenth century[21].

Busra al-Harir contains 'extensive ancient and massive' ruins with Greek inscriptions[22]. In the sixteenth century Busra al-Harir was an entirely Muslim settlement with a population of 45 households and 35 single men[23]. Najran, however was exclusively Christian up to the end of the seventeenth century when Druze fleeing persecution in Mount Lebanon settled in the village[24] (Firro 1992, 38-39 and Map No.4). The village contains many ruins and ancient buildings dominated by a sixth century church with two towers[25]. The village of Harran also contains extensive early remains, the most significant of which is a bilingual Arabic Greek inscription dated to 568 AD[26].

**Bedouin Burials**

The Tell Qarassa graves differ from most other excavated examples of early Islamic burials because they were not located in a cemetery and – from the available evidence- do not seem to have been located near a permanent settlement of the period. Based on this information and also taking into account the DNA results it seems likely that the deceased were Bedouin. Whilst Bedouin are sometimes regarded as the autochthonous form of Arab, in practice they are often regarded as marginal and as such may not have qualified for burial in one of the nearly settlements. The Leja region has a long history of occupation by Bedouin nomads and it is known that the area of Tell Qarassa was occupied by Bedouin of the Banu Sarma in the sixteenth century[21].

Because of the long-term continuity of the Bedouin nomadic lifestyle, burial practices of today or the recent past are regarded as comparable with the distant past back to the early years of Islam[27]. There are a number of aspects of the Tell Qarassa burials which can be documented in Bedouin burials both from the recent and more distant past. Important studies discussing Bedouin burial practices include William and Fidelity Lancaster's (1993) study of the Bedouin graves in north-eastern Jordan[28] and Mustafa and Abu Tayeh's (2013) study of Bedouin funerary practices[29] based on comparing western travellers accounts, comments from focus groups and personal observation. Mustafa and Abu Tayeh[29] draw attention to the fact that many nineteenth century European travellers were unaware of the differences between sedentarized and mobile nomadic Bedouin and thus confused many details of village customs with those of the mobile tribes.

One of the most obvious features of the Tell Qarassa burials is that there were just two graves and no communal cemetery. From studies of nomadic Bedouin, it appears that when someone dies, they are buried immediately in a prominent nearby location[29]. There is a concept of tribal burial grounds located at places with particular associations such as the tomb of the founder of a tribe. In this case the tribe concerned will make an effort to bury the deceased in this location and amongst the Bedouin of Sinai there is a tradition of carrying bodies for several days prior to burial. Mustafa and Abu Tayeh[29] also draw attention to the changing dynamics of nomad groups thus individual tribes or tribal groupings will move spatially through time thus a Bedouin tribe resident in Syria today may have been located in Jordan, Saudi Arabia or even Iraq in early Islamic times. In this respect it is worth noting the genetic evidence from Israel cited in the main text[30] which suggests that the Negev Bedouin arrived around 700 AD.

**Re-Use of Ancient Sites**

The precise choice of burial ground often involves the re-use of an ancient site; famous examples in the Negev include Tell el-Hesi, Arad, Tell Sheva, Tell Malhatta and Arad[31]. In Syria there are also many examples and at Tell Abu Hureyra, a late Islamic cemetery which covered the Neolithic site, was fully excavated[32]. In some cases the re-use of an ancient site can cause problems of interpretation thus some of the burials from the Qumran cemetery in Palestine/Israel which were previously thought to belong to the Essene community of the 3rd century BC to 1st century AD have been re-interpreted as belonging to Bedouin from the Mamluk (1260-1516 AD) or Ottoman period (1516-1918)[31]. There are a number of reasons for the re-use of ancient sites as burial grounds - some of these are conscious whilst others may be unconscious. Ancient sites, and in particular Tells are usually higher than the surrounding landscape and form both an important focal point and a viewing platform. Other advantages of ancient sites include the fact that they often have readily available stone which can be used to form a cairn on top of the grave both to deter animals from digging up the body and also to mark the position of the grave. The reality of the threat posed by animals digging up recently buried people is proved by finds of human bones within hyena dens in the Judean desert[31,33]. Also, the deceased persons' clothes are sometimes placed on top of the grave and held down with stones[29]. Another practical reason why ancient sites may be chosen as burial locations is that they are not usually used for cultivation or other agricultural activities which may disturb the graves.

There may, however, be fewer practical reasons for the choice of an ancient site as a burial ground. According to Muslim tradition after his death Muhammad was buried within the confines of his house in Medina[34]. This practice is still sometimes carried out by semi-nomadic Bedouin and examples can be found in the Negev today. In any case derelict buildings were often regarded as good burial locations and Johann Burckhardt noted that if a Bedouin camp is located near an abandoned village and one of the members of the tribe dies the preference is for them to be buried in the ruin[35]. Also ruins have a special status within pre-Islamic Arabic poetry which frequently refer to ruins as symbols of loss and the passing of time. Thus, the pre-Islamic poet Imru al-Qais expressed his sorrow in the following words:

Stop, Oh my friends, let us pause to weep over the remembrance of my beloved.

Here was her abode on the edge of the sandy desert between Dakhool and Howmal.

The traces of her encampment are not wholly obliterated even now.

For when the South wind blows the sand over them the North wind sweeps it away.

(*Mu'allaqat* Imru al-Qais[36]).

In most cases where ruins are re-utilized as burial sites there is a long gap between the domestic use of a location and its re-use as a burial site. This suggests that those responsible for the burial will have no knowledge or connections with the site although when digging a grave they will be aware of the antiquity of the site particularly if there are earlier pre-historic burials present. This question has been discussed in relation to Çatal Höyük where it has been suggested that there is an awareness of the deep past from the almost continuous use of the site as a burial ground from pre-history to the twentieth century[37]. However, this situation is unusual and, in any case, involves sedentarised villagers rather than nomadic Bedouin.

One other way in which ancient sites may be of relevance to the Bedouin is the concept of land ownership. By burying members of the tribe at ancient locations the Bedouin may be asserting their rights to camp in the area. The ancient site provides a form of authority whilst the burial of a tribal member forms a connection with that authority.

**Identity of the Deceased**

The close proximity of the two graves suggests that two individuals were related and may have been cousins. Given the proximity of the $^{14}C$ dates and the fact that these are the only non-prehistoric burials at the site, it seems likely that they died either at the same time or shortly one after the other. The absence of trauma to the bones and the young age of the deceased suggests that they may have died from a disease, possibly the Justinian plague which ravaged the Middle East from 541 AD to 749 AD recurring in cycles of nine to twelve years[38,39]. Specifically, the dates of the burials may be linked to the outbreak of 79 AH (698 AD) which was reported in Syria by as-Suyuṭī[39].

The specific identity of the two people buried at Tell Qarassa in the late seventh or early eighth century as revealed by the DNA analysis provides a tantalizing glimpse of early Islamic society in Syria. At present there are no examples of DNA studies from the region which relate to this period and worldwide the only DNA analysis which relates to early Islamic burials is the study of two individuals from the south of France[4].

Within the context of early Islamic and Late Antique Syria discussed above, the two individuals from Tell Qarassa could fit either of the two scenarios suggested by genome sequencing. In the first scenario the individuals are representative of a Bedouin Arab society already present in Syria before the Muslim conquest whilst the second scenario suggests that both people were the children or grandchildren of a family which had migrated to Syria with the Muslim conquest. Whilst it is not possible to identify which of the two scenarios applies to these individuals this region of southern Syria had direct connections with Arabia and in particular the Hijaz for centuries[40]. In pre-Islamic times these connections were in the form of direct trade between Mecca and the Hauran. We know for example that one of Muhammad's ancestors Hāshīm organized a bi-annual caravan carrying grain from the Hauran to Mecca and that as a young man Muhammad travelled to Bosra as a merchant (Ṭabarī cited in[40]). The direct connection between Mecca and the Hauran was continued in Islamic times with the annual pilgrimage or Hajj. In addition to the passage of pilgrims the Hauran continued to send supplies to Mecca. The nature of material traded from Mecca to Syria has been a matter of academic controversy, however it seems likely that materials from southern Arabia were probably traded via Mecca as was coffee during the sixteenth and seventeenth centuries[41].

We also know from Late Antique inscriptions in Bosra and the Hauran region that there were substantial family connections with Arabia attested in Nabatean and Safaitic names. Maurice Sartre states that the Quraysh from Mecca probably had agents living in Bosra[40]. Turning from genetic to religious identity, the fact that both individuals were buried according to Muslim tradition is very interesting. Firstly, it provides additional evidence for the early adoption of specific Islamic burial rites which were followed even in remote locations. Secondly it is worth pointing out that although the Hauran was incorporated into the territory of Islam (*Dar al-Islam*) from an early date much of the population remained Christian. The enduring strength of Christianity in the region is proved by the fact that two churches were built several decades after the Muslim conquest one in 652 (Kafr) and one in 668 ('Orman)[40]. It therefore seems likely that the two individuals buried at Tell Qarassa were not part of the local sedentarised Arab society which was predominantly Christian but were instead Muslims who were far from the nearest Islamic cemetery. Their presence in the area may either have been because they were nomads camping in the area or perhaps, they were pilgrims utilising one of the near-by Hajj (pilgrim) routes which connected the region to Mecca.

**Significance**

This study provides further insight into the re-use of prehistoric sites as cemeteries by Muslims and in particular Bedouin. This example raises the question of whether the people undertaking the burial were in some way conscious of the deep history of the site or were simply using it as a convenient location. Given the presence of Neolithic graves on the site, it seems probable that those digging the graves in the late seventh or early eighth centuries AD may have been aware of the earlier internments. Did the presence of these earlier graves make it more acceptable for the early Muslims to bury their relatives in this remote area?

The DNA analysis of the two graves from Tell Qarassa is only the second time that genome sequencing has been applied to early Islamic burials and as such is of considerable significance in efforts to understand the human origins of this dynamic cultural period. The first project to utilise DNA evidence focused on three seventh to ninth century Islamic burials from Nimes in France. The analysis showed that all three individuals were males of Berber and/or Arab and Berber origin and as such provided a useful supplement to the patchy historical information about the period and also the first evidence for a Muslim presence in France[4]. Similarly, the present study provides definitive proof of the early presence of Muslim Arabs in the Syrian countryside and provides more evidence of the Bedouin contribution to early Islamic society.

### Supplementary Text S2

#### *Additional unsupervised ADMIXTURE runs to verify the choice of sources*

In the unsupervised ADMIXTURE that was run with these ancient populations (Fig. S5), we observed that African sources (Hadza, Mbuti and Yoruba) were immediately differentiated from other groups (K = 2). At K = 3, a new component emerged in Levant_N, Anatolia_N, Bedouins (almost entirely in Bedouin B), Saudis and a few Middle Eastern populations. At K = 4, African sources are further differentiated and at K = 5, WHG are seen to form a separate cluster with the corresponding new component also emerging in Levant_N, Levant_BA, Anatolia_N, Iran_Chl, Turkish and Jewish groups and our samples. At K = 6, Mbuti and Taa_West are differentiated from Yoruba. At K = 7, a new component appears in Anatolia_N, and a major fraction is also seen in Levant_N, Levant_BA and to a lesser extent in Iran_Chl and our samples. At this value of K, 9/10 iterations were found to agree and K = 7 was chosen as the best fit. Between our samples and the Bedouin B, differentiation appears at K = 5, where the component that separates out WHG is seen in our samples and then at K = 7, where this disappears and a small proportion of Anatolia_N component emerges. Overall, we observed that our samples clustered most similarly with Levant_N and Levant_BA. Small proportions of WHG were seen at K = 5, 6 and a very small proportion of the component (pink) that constitutes the majority of Iran_N is observed as well. This was also true for Levant_BA and Levant_N, and thus overall corresponds to the supervised admixture results.

### Supplementary Figures

Fig S1. (a) Plots of read length distribution; (b) Patterns of base misincorporation at fragment termini.

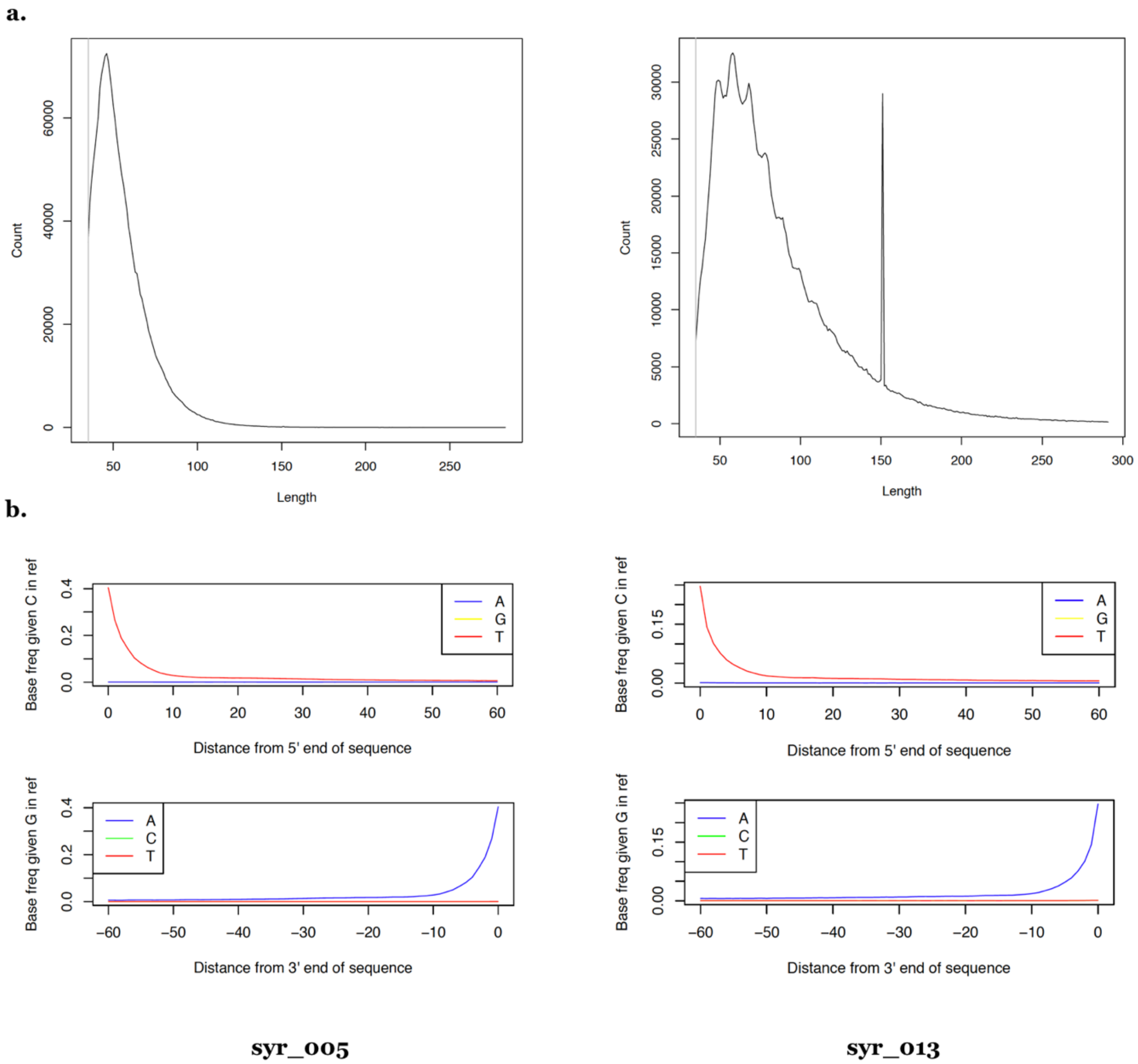

Fig S2. Results of unsupervised ADMIXTURE for K = 2 to K = 5

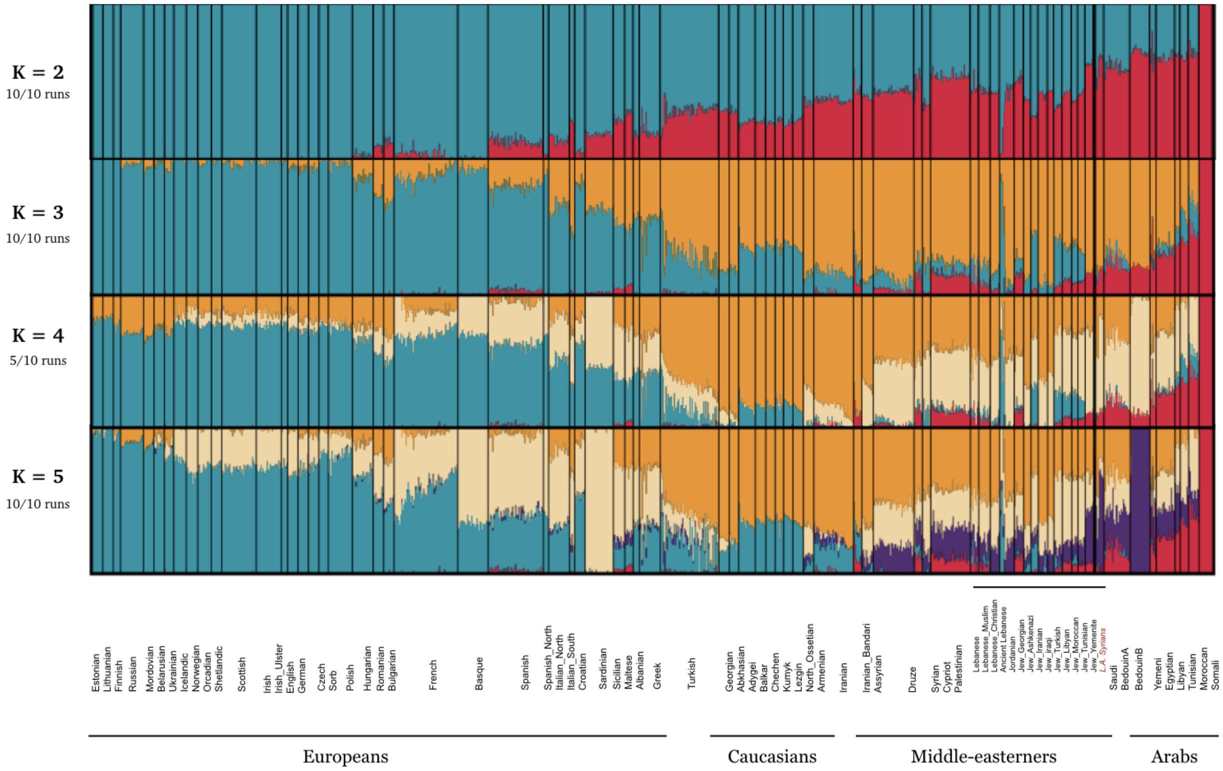

Fig S3. Supervised ADMIXTURE proportions.

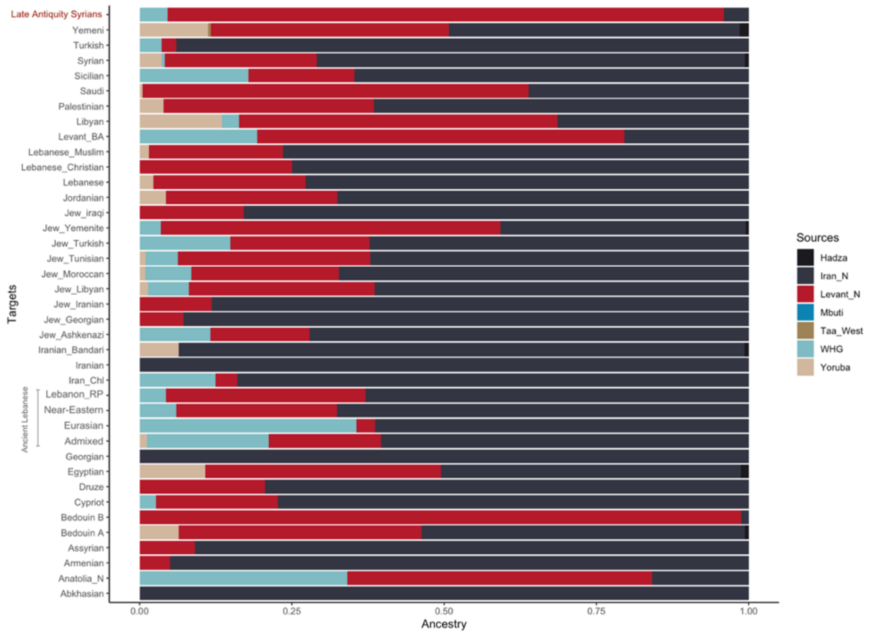

Figure S4. CND estimates

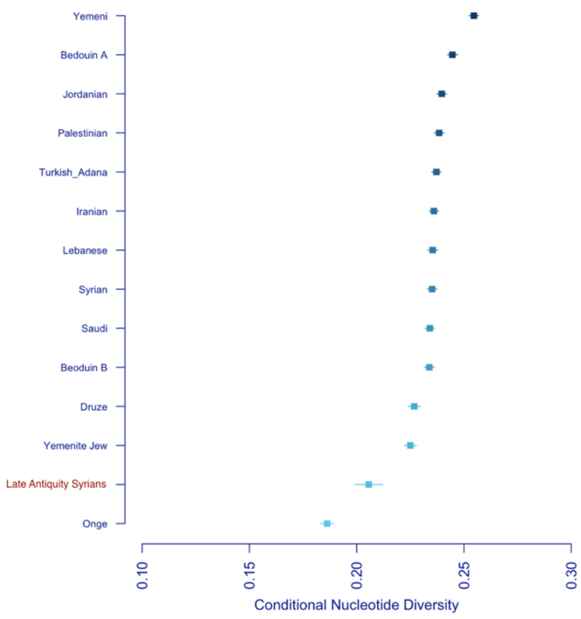

Figure S5. Results of unsupervised ADMIXTURE runs to verify the choice of ancient populations for supervised ADMIXTURE.

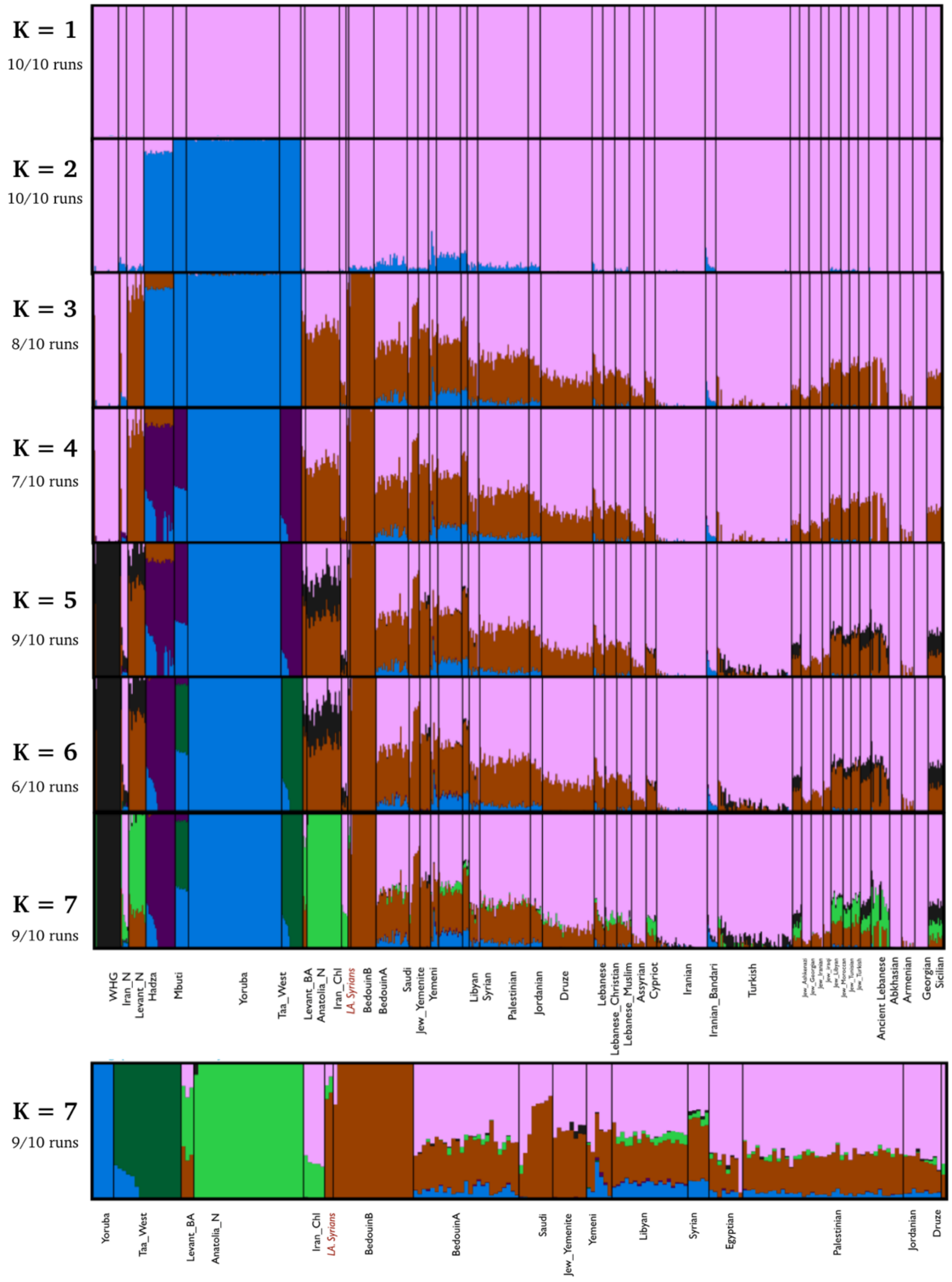

### Supplementary Tables

*Table S1. Estimates for mitochondrial, X-chromosomal and autosomal contamination.*

| Sample | Genome Cov. | Mt Cov | Mt Point estimate | Authentic DNA | X chromosomal | Autosomal |
|---|---|---|---|---|---|---|
| syr005 | 0.159799 | 23.5158 | 0 | 0.9974079 | 0.027 ± 0.015 | 0.01475 |
| syr013 | 6.15177 | 522.349 | 0.2 | 0.9997392 | NA | 0.0488 |

*Table S2. List of derived Y-chromosome markers for syr005.*

| Marker | Haplogroup | State |
|---|---|---|
| L1013 | A1b | derived |
| PF985 | BT | derived |
| M9129 | BT | derived |
| Z17372 | BT | derived |
| M9344 | BT | derived |
| Z40409 | BT | derived |
| M11760 | BT | derived |
| M5758 | CT | derived |
| Z17706 | CT | derived |
| PF3553 | IJ | derived |
| PF4595 | J | derived |

*Table S3. Values of Outgroup f3 statistics calculated between Syrian samples and Middle-east Groups*

| Population | Outgroup f3 values | Standard Error | Z-score |
|---|---|---|---|
| Abkhasian | 0.254594855789 | 0.00318325880687 | 79.9793140411 |
| Adygei | 0.25261711279 | 0.00312097736488 | 80.9416677073 |
| Armenian | 0.256306101585 | 0.00312184922799 | 82.1007303259 |
| Balkar | 0.24991321127 | 0.00308256981459 | 81.0730093076 |
| Chechen | 0.25409044703 | 0.00319580365215 | 79.507527585 |
| Georgian | 0.254876365344 | 0.00316822594601 | 80.4476605165 |
| Kumyk | 0.25061602306 | 0.00318127729056 | 78.7784277101 |
| Lezgin | 0.25381576998 | 0.00314674866054 | 80.6596895273 |
| North_Ossetian | 0.251276142262 | 0.00308113163712 | 81.5531992318 |
| Jew_Ashkenazi | 0.253817916158 | 0.00323394694435 | 78.4854917307 |
| Jew_Georgian | 0.257439285657 | 0.0032554989102 | 79.0782896135 |
| Jew_Iranian | 0.255510985398 | 0.00313014924082 | 81.6290105489 |
| Jew_iraqi | 0.257502031096 | 0.00322457984111 | 79.8559948224 |
| Jew_Libyan | 0.251014346027 | 0.00307274610741 | 81.690558625 |
| Jew_Moroccan | 0.250811904783 | 0.00318326435314 | 78.7907873675 |
| Jew_Tunisian | 0.251031739979 | 0.00315785663094 | 79.4943435743 |
| Jew_Turkish | 0.253550140602 | 0.00310015310217 | 81.7863286897 |
| Jew_Yemenite | 0.254212744532 | 0.00320208510493 | 79.3897526773 |
| Turkish | 0.252926010357 | 0.00305120903019 | 82.8937014324 |
| Assyrian | 0.256787392083 | 0.00313905014796 | 81.8041700449 |
| Syrian | 0.2487563738 | 0.00302800469242 | 82.1519115947 |
| Cypriot | 0.258925502145 | 0.00318565300986 | 81.2786268132 |
| Druze | 0.254734125606 | 0.00309133611239 | 82.4025975645 |
| Jordanian | 0.246316466428 | 0.00306156552561 | 80.4544160062 |
| Palestinian | 0.24762307819 | 0.00301120610035 | 82.2338524622 |
| Lebanese_Christian | 0.256105828128 | 0.00309131694565 | 82.8468360349 |

| | | | |
|---|---|---|---|
| Lebanese | 0.250572009629 | 0.00309138325109 | 81.0549806596 |
| Lebanese_Muslim | 0.253559637531 | 0.00313785442783 | 80.8066923955 |
| Iranian_Bandari | 0.237813908478 | 0.00302638772186 | 78.5801193814 |
| Iranian | 0.25055504783 | 0.00305621971934 | 81.9820140038 |
| **Saudi** | **0.261020315354** | **0.00318728557576** | **81.8942354395** |
| BedouinA | 0.243890411538 | 0.00296593967958 | 82.2304017905 |
| **BedouinB** | **0.260777577371** | **0.0030912182945** | **84.3607770552** |
| Yemeni | 0.233549814908 | 0.00306805239234 | 76.1231507944 |
| Libyan | 0.234981498863 | 0.00307654572193 | 76.3783542003 |
| Egyptian | 0.235740590839 | 0.00292750777152 | 80.5260341687 |
| Sicilian | 0.25402133036 | 0.00309781843235 | 82.0000706649 |

*Table S4a. Top 25 values of D-statistics calculated for the topology (Syr, Bedouin A: X, Mbuti)*

| **Population *X*** | **D-Statistic** | **Standard Error** | **Z-score** |
|---|---|---|---|
| BedouinB | 0.0564629628856 | 0.00410478341373 | 13.7554061188 |
| Saudi | 0.0549399728896 | 0.00409865798404 | 13.4043809226 |
| Ancient Lebanese | 0.0412605758856 | 0.00425000330273 | 9.70836325213 |
| Jew_Yemenite | 0.0409971343714 | 0.0041493478683 | 9.88038016399 |
| Libyan | 0.0390960272233 | 0.00407663947135 | 9.5902587163 |
| Yemeni | 0.038057253172 | 0.00403666699855 | 9.42789018408 |
| Jew_iraqi | 0.0378467658643 | 0.00422311149258 | 8.96182019602 |
| Lebanese | 0.0375242122338 | 0.00393420392806 | 9.53794285197 |
| Syrian | 0.0371670760501 | 0.00384304726369 | 9.67125135341 |
| Cypriot | 0.0366696596752 | 0.00411742400429 | 8.90597121816 |
| Assyrian | 0.0364723548016 | 0.00405615882021 | 8.99184583697 |
| Moroccan | 0.0362499922496 | 0.00359683978012 | 10.0782894056 |
| Lebanese_Muslim | 0.036164081469 | 0.00397240201047 | 9.10383223391 |
| Palestinian | 0.035597688354 | 0.00379190217237 | 9.38781823366 |
| Jordanian | 0.0355131895665 | 0.00387181472379 | 9.17223371982 |
| Druze | 0.0349766854731 | 0.00388646113716 | 8.99962311181 |
| Egyptian | 0.0349129648335 | 0.00371808945157 | 9.39002820892 |
| Tunisian | 0.0348950173804 | 0.00375081156456 | 9.30332456849 |
| Jew_Libyan | 0.0345635848521 | 0.00404495795538 | 8.5448563949 |
| Jew_Georgian | 0.0342782325103 | 0.0042269529855 | 8.1094425767 |
| Lebanese_Christian | 0.0340240847486 | 0.00404028181464 | 8.42121572444 |
| Sardinian | 0.0338321471761 | 0.00406666409122 | 8.31938572186 |
| Jew_Iranian | 0.033565728223 | 0.00409277533107 | 8.2012144591 |
| Spanish_North | 0.0334249294099 | 0.00428205976085 | 7.80580638213 |

*Table S4b. Top 25 values of D-statistics calculated for the topology (Syr, Bedouin B: X, Mbuti)*

| **Population *X*** | **D-Statistic** | **Standard Error** | **Z-score** |
|---|---|---|---|
| BedouinB | -0.0341610012937 | 0.0039782336333 | -8.58697714677 |
| Ancient Lebanese | 0.026291560906 | 0.00441379000542 | 5.95668594873 |
| Saudi | 0.0261054460408 | 0.00409937098695 | 6.36815895023 |
| Jew_iraqi | 0.022295642107 | 0.0042744515017 | 5.216024114 |
| Assyrian | 0.0221829651713 | 0.00418200179668 | 5.30438920158 |
| Lebanese | 0.0219825434304 | 0.00400912646252 | 5.48312547281 |
| Cypriot | 0.0217910764549 | 0.00413966754434 | 5.2639677514 |
| Moroccan | 0.0213703151225 | 0.00368880322283 | 5.79329224996 |
| Syrian | 0.0208263403396 | 0.00388480243161 | 5.36097799213 |
| German | 0.0207697146259 | 0.0042561278658 | 4.87995550904 |
| Jew_Georgian | 0.0202655849564 | 0.00423592624479 | 4.78421572645 |
| Spanish_North | 0.0202254379988 | 0.00440115353719 | 4.59548566708 |
| Sardinian | 0.0200991237333 | 0.00409252836798 | 4.91117517731 |
| Jordanian | 0.0200382229666 | 0.00391563563429 | 5.1174891737 |
| Albanian | 0.0199370934282 | 0.00430281978698 | 4.63349487434 |
| Sorb | 0.019936714119 | 0.00423948875342 | 4.70262224494 |
| Lezgin | 0.0197530964642 | 0.00410136374135 | 4.81622643343 |
| Jew_Libyan | 0.0197093231216 | 0.00409031639548 | 4.81853265516 |

| | | | |
|---|---|---|---|
| Maltese | 0.0197011520089 | 0.00417564163284 | 4.71811370353 |
| Lebanese_Muslim | 0.0196879048127 | 0.00400301053448 | 4.91827454441 |
| Finnish | 0.019646681016 | 0.00426512937165 | 4.60635054745 |
| Hungarian | 0.0196343047155 | 0.00406343249462 | 4.83195051018 |
| Tunisian | 0.0195532050442 | 0.00379217564927 | 5.15619708912 |
| English | 0.019514087367 | 0.0041218570744 | 4.73429500703 |

##### *Table S4c. Top 25 values of D-statistics calculated for the topology (Syr, Saudi: X, Mbuti)*

| **Population *X*** | **D-Statistic** | **Standard Error** | **Z-score** |
|---|---|---|---|
| Saudi | -0.0583929348666 | 0.00405681047637 | -14.3938040011 |
| BedouinB | 0.0255641389308 | 0.00424494947752 | 6.02224810121 |
| Ancient Lebanese | 0.01929440332 | 0.00446912820634 | 4.31726332949 |
| Moroccan | 0.0176625967744 | 0.00373052650449 | 4.73461232701 |
| BedouinA | 0.0174323214495 | 0.00388842650291 | 4.48312998496 |
| Jew_iraqi | 0.0158841076468 | 0.00434472918863 | 3.65594884218 |
| Cypriot | 0.0156608350011 | 0.00421086226633 | 3.71915156816 |
| Libyan | 0.0156206341263 | 0.00425416414356 | 3.67184565503 |
| Jew_Yemenite | 0.0151611194623 | 0.00434006761904 | 3.49329107126 |
| Assyrian | 0.0149357199408 | 0.00425466316801 | 3.51043533907 |
| Tunisian | 0.0147813862816 | 0.00395616907018 | 3.73628781261 |
| Lebanese_Muslim | 0.0147723022293 | 0.00414361377426 | 3.56507701588 |
| Lebanese | 0.0145790673824 | 0.00406060071479 | 3.59037206718 |
| German | 0.014250387461 | 0.00430932890888 | 3.30686929736 |
| Palestinian | 0.0140957384175 | 0.00392958122517 | 3.58708412165 |
| Jew_Libyan | 0.0140370839426 | 0.00418271911731 | 3.35597097222 |
| Yemeni | 0.0139012880884 | 0.0042479045251 | 3.27250483298 |
| Syrian | 0.0138193977134 | 0.00400560963958 | 3.45001109865 |
| Sardinian | 0.0137307871554 | 0.00417695368034 | 3.28727302388 |
| Jordanian | 0.0135778386864 | 0.00406052461945 | 3.34386316029 |
| Egyptian | 0.0135603758226 | 0.00388708016814 | 3.48857631847 |
| Jew_Georgian | 0.0135152209864 | 0.00431782824441 | 3.13009694258 |
| Druze | 0.0132958064824 | 0.00403972788262 | 3.29126289411 |
| Maltese | 0.0130947512882 | 0.00426635656552 | 3.06930541016 |

##### *Table S4d. Top 25 values of D-statistics calculated for the topology (Syr, Jew_Yemenite: X, Mbuti)*

| **Population *X*** | **D-Statistic** | **Standard Error** | **Z-score** |
|---|---|---|---|
| Jew_Yemenite | -0.0753024146011 | 0.00410592659086 | -18.3399320311 |
| BedouinB | 0.0335927527277 | 0.00424168923221 | 7.9196638152 |
| Saudi | 0.0302876568828 | 0.00424964696151 | 7.12709953488 |
| Ancient Lebanese | 0.0208475673664 | 0.00444888240869 | 4.6860234664 |
| Moroccan | 0.0193906723602 | 0.00366894165063 | 5.2850860566 |
| Libyan | 0.0190751952451 | 0.00424595734584 | 4.49255460932 |
| BedouinA | 0.0183528406453 | 0.00382403887591 | 4.7993342225 |
| Syrian | 0.0171224631778 | 0.00392302821415 | 4.3646036284 |
| Tunisian | 0.0168021251074 | 0.00393390958337 | 4.2711009878 |
| Lebanese | 0.0166019772202 | 0.00407245172731 | 4.0766541464 |
| Jew_iraqi | 0.0155464166948 | 0.00431478166375 | 3.60305987795 |
| Assyrian | 0.0154872120216 | 0.0042554649892 | 3.63937009489 |
| Jew_Libyan | 0.0154399362738 | 0.00413479798519 | 3.73414525428 |
| Lebanese_Muslim | 0.0152968421969 | 0.0041097853328 | 3.72205381988 |
| Yemeni | 0.0152780055946 | 0.00414693654087 | 3.68416672019 |
| German | 0.0151053152765 | 0.00431762326237 | 3.49852554486 |
| Jordanian | 0.0148170450013 | 0.00404032788569 | 3.66728775992 |
| Spanish_North | 0.0147878738893 | 0.00445694575626 | 3.31793894251 |
| Finnish | 0.0147547347438 | 0.00428253797568 | 3.4453249049 |
| Egyptian | 0.0145873888305 | 0.00385872929519 | 3.78036076506 |
| Albanian | 0.0144624820943 | 0.00444636406934 | 3.25265359939 |

| | | | |
|---|---|---|---|
| Druze | 0.0143674119715 | 0.00403675511456 | 3.55914876275 |
| Icelandic | 0.0143296074252 | 0.00420677077049 | 3.40632000339 |
| Cypriot | 0.014289529261 | 0.00420532946061 | 3.39795713864 |

*Table S5. Values of Conditional Nucleotide Diversity calculated for various Middle-east groups*

| Sites Differing | Sites Covered | CND Estimate | Standard Error | Samples Compared |
|---|---|---|---|---|
| 140044 | 598861 | 0.233850593042 | 0.00121061614884 | BedB_607-608 |
| 146980 | 600819 | 0.244632742973 | 0.00114811333385 | BedA_609-611 |
| 143174 | 597388 | 0.23966668229 | 0.00115364667638 | Jordan_214-307 |
| 142137 | 596114 | 0.238439291813 | 0.00116059735487 | Palestinian_675-676 |
| 140107 | 598468 | 0.234109426068 | 0.00113215608327 | Saudi_A5-A6 |
| 141431 | 599294 | 0.235996021986 | 0.0011065770667 | Iran_11-14 |
| 134216 | 596507 | 0.225003227121 | 0.00140128791805 | Yemenite_Jew_4684-95 |
| 135894 | 599073 | 0.226840468524 | 0.00140013283536 | Druze_557-558 |
| 141112 | 599423 | 0.235413055555 | 0.00117347124191 | Lebanon_1-2 |
| 142524 | 600990 | 0.237148704637 | 0.00111169782973 | TurkAdana_23108-112 |
| 109298 | 586917 | 0.186223946486 | 0.00159632020817 | Onge1-12 |
| **4024** | **19571** | **0.205610341832** | **0.0032747874575** | **Syr_005-013** |
| 152487 | 598901 | 0.254611363147 | 0.00110476097775 | Yemen2-Yemen3 |
| 140958 | 99437 | 0.235150649693 | 0.0011392422505 | syria364-syria4 |

*Table S6. Lactase persistence variants tested in the samples. The number of reads mapping in syr013 are shown, as well as the alleles seen at the variant site in the reads.*

| SNP ID | Chr | Position (GRCh37) | Ancestral | Mutation | Referred as | No. of Reads Mapped | Alleles Observed | Region of Prevalence | Remarks |
|---|---|---|---|---|---|---|---|---|---|
| rs4988235 | 2 | 136608646 | G | G > A; G > C | -13,910*T | 10 | GAGGGGGGGG | Europe | Allele A associated. Also found in some populations from Africa, South Asia |
| **rs41380347** | 2 | 136608651 | A | A > C; A > G | -13,915*G | 9 | AACCCCCAA | Arabian Peninsula | Allele C associated. East and North Africa, Middle East |
| rs145946881 | 2 | 136608746 | C | C > T; C > G | -14,010*C | 10 | CCCCCCCCCC | Africa | Allele G associated. Identified in East Africa |
| rs41525747 | 2 | 136608643 | G | G > C | -13,907*G | 10 | GGGGGGGGGG | Africa | Allele C associated. Sudan and Ethiopia |
| rs182549 | 2 | 136616754 | C | C > T | G/ A*14107 | 11 | CCCCCCCCCCC | Africa | Allele T associated. Rare variant, Xhosa |

*Table S7. Phenotypic variants tested in the samples. The clinical significance (as described in dbSNP) of the variant alleles is shown, along with additional remarks about the inheritance/region of prevalence.*

| Condition | SNP ID | Chr | Position (GRCh37) | Ancestral | Mutation | Reads Mapping | Reads | Remarks |
|---|---|---|---|---|---|---|---|---|
| Familial hypercholesterolemia | rs5742904 | 2 | 21229160 | C | C > A, C > T | 8 | CCCCCCCC | Allele A pathogenic; T likely pathogenic. Autosomal Dominant |
| G6PDH Deficiency | rs5030868 | X | 153762634 | G | G > A | 5 | GGGGG | Other/Likely pathogenic. |
| G6PDH Deficiency | rs1050828 | X | 153764217 | C | C > T | 6 | CCCCCC | Allele T pathogenic. 'Middle-east' variant |
| Sickle-cell anemia | rs334 | 11 | 5248232 | T | T > A, C, G | 5 | TTTTT | Allele A other/ pathogenic; G other. |
| Bardet-Biedl Syndrome | rs28938468 | 15 | 73027508 | C | C > A, G | 6 | CCCCCC | Allele A pathogenic. Autosomal Recessive |
| Bardet-Biedl Syndrome | rs28937875 | 20 | 10394008 | C | C > T | 8 | CCCCCCCC | Allele T pathogenic. Pleiotropic Disorder |
| Bardet-Biedl Syndrome | rs2277598 | 15 | 73027478 | T | T > C | 8 | CCCCCCC^FC | Allele C benign/minor contributor |
| Meckel-Gruber Syndrome | rs386834180 | 8 | 94793953 | T | T > C | 4 | TTTT | Allele C likely pathogenic. Autosomal Recessive, pleiotropic and lethal |
| Osteopetrosis | rs398123011 | 7 | 26404195 | G | G > A | 8 | GGGGGGGG | Allele A pathogenic. Autosomal Recessive |
| Marfan Syndrome | rs113871094 | 15 | 48758017 | G | G > A | 6 | GGG$GGG | Allele A pathogenic. Autosomal Dominant |
| Marfan Syndrome | rs137854468 | 15 | 48779593 | C | C > T | 7 | CCCCCCC | Allele T pathogenic. Autosomal Dominant. |
| Congenital chloride diarrhea | rs386833481 | 7 | 107431671 | G | G > A, C | 7 | GGGGGGG | Allele A likely pathogenic; C: uncertain significance. Autosomal Recessive |
| Hypophosphatemic rickets | rs121908249 | 6 | 132211575 | A | A > C, G | 10 | AAAAAAAAAA | Allele C pathogenic. Autosomal recessive, prevalent in Bedouins[42] |

*Table S8. Collagen stable isotope data for the samples analysed in this paper, with similar data for humans and fauna from the Middle East from proximate periods.*

| **Sample Code** | **Location** | **Species** | **Collagen yield(%)** | **C%** | **N%** | **C:N** | **$\delta^{13}C$** | **s.d. ($\delta^{13}C$)** | **$\delta^{15}N$** | **s.d. ($\delta^{15}N$)** | **Age** | **Reference** |
|---|---|---|---|---|---|---|---|---|---|---|---|---|
| Syr 005 | Syria | Human | 13.6 | 42.25 | 15.26 | 3.23 | -19.2 | | 13.1 | | Late Antique | This paper |
| Syr 013 | Syria | Human | 17.5 | 40.98 | 14.79 | 3.24 | -18.5 | | 11.5 | | Late Antique | This paper |
| LR/Byz Humans | Jordan | Human (n=7) | N/A | N/A | N/A | N/A | -19.1 | 0.3 | 8.1 | 0.6 | Late Roman/Byz | Sandias & Müldner (2015)[43] |
| LR/Byz Sheep/goats | Jordan | Sheep/ goats (n=5) | N/A | N/A | N/A | N/A | -19.3 | 0.5 | 5.3 | 0.9 | Late Roman/Byz | Sandias & Müldner (2015)[43] |
| LR/Byz cattle | Jordan | cattle (n=2) | N/A | N/A | N/A | N/A | -18.8 to -15.6 | | 5.4 to 9.7 | | Late Roman/Byz | Sandias & Müldner (2015)[43] |
| PAR | Syria | Human (n=2) | N/A | N/A | N/A | N/A | -19.5 to -19.1 | | 9-9.3 | | | Sołtysiak & Schutkowski (2015)[44] |
| MOD | Syria | Human (n=4) | N/A | N/A | N/A | N/A | -18.9 | 0.5 | 8.8 | 0.5 | | Sołtysiak & Schutkowski (2015)[44] |
